# A selective and augmentable FFAR2 signal circuitry programs a butyrate-induced cellular identity of enteroendocrine L-cells

**DOI:** 10.1101/2025.06.14.659710

**Authors:** Aanya Hirdaramani, Chia-Wei Cheng, Aylin C. Hanyaloglu, Gary Frost

## Abstract

Activation of free fatty acid receptor 2 (FFAR2) on enteroendocrine L-cells mediates secretion of glucagon-like peptide 1 (GLP-1) and peptide YY (PYY), key regulators of central appetite control with therapeutic relevance to obesity. Here, we show that butyrate, a metabolite derived from fermentation of dietary fibre and an FFAR2 agonist, stimulates a PYY-biased profile in a human L-cell model at the transcriptional, morphological and secretory level via an FFAR2-Gαi axis that does not require dynamin-dependent receptor internalization. We observe that butyrate modulates active Notch cascades within a Hes1-GFP mouse organoid model, which are antagonistic to secretory differentiation, and identify butyrate-dependent regulation of late-stage human enteroendocrine maturation markers, *NeuroD1* and *Pax6*. Butyrate-mediated upregulation of *Pyy* and *Pax6* is enhanced by the FFAR2-selective Gαi biased allosteric agonist AZ-1729. Our study reveals functions of spatiotemporally-regulated butyrate-activated FFAR2 signalling mechanisms that could be pharmacologically amplified to fine-tune L-cell populations in the human colon.

**Graphical Abstract:** 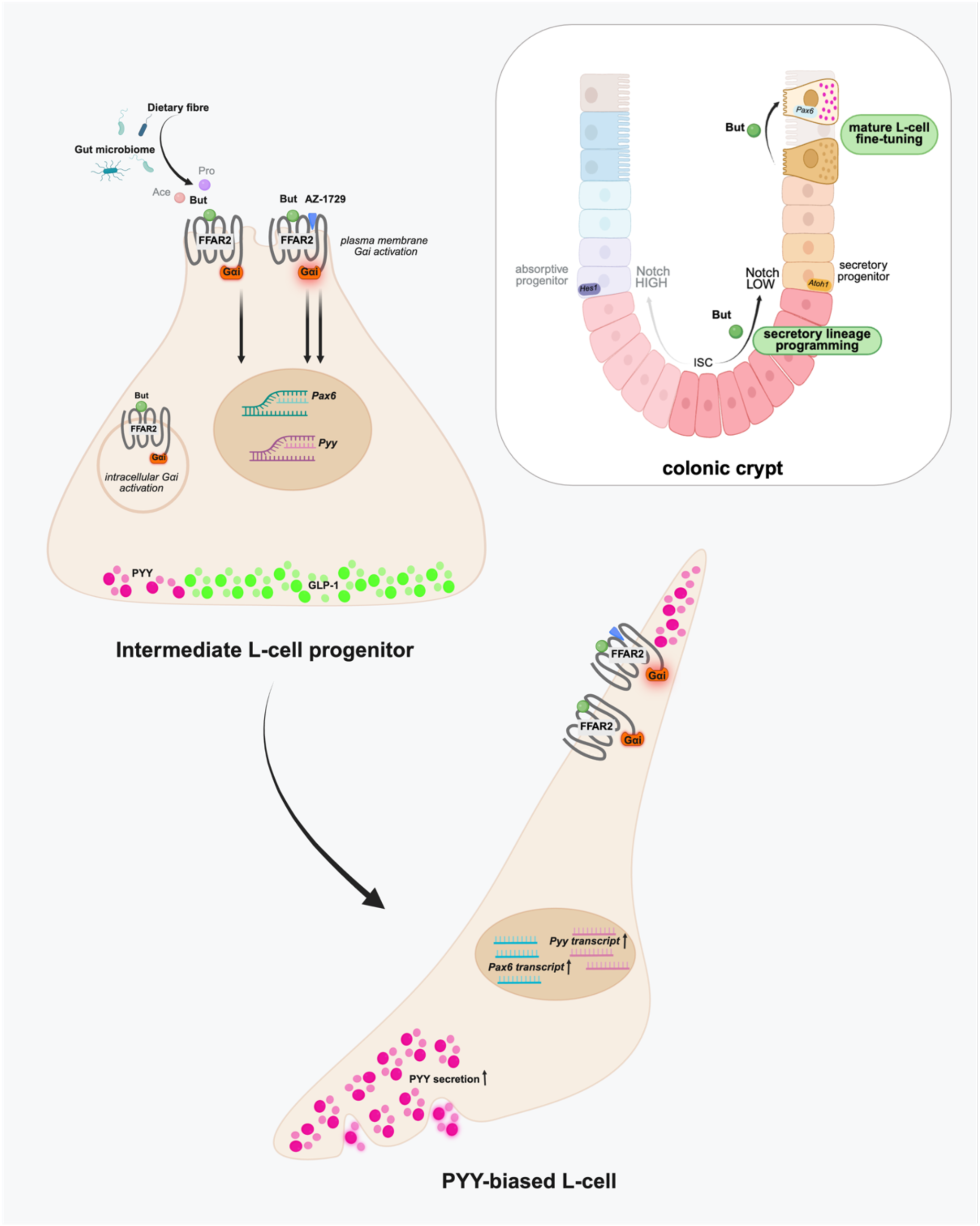

## Introduction

Increased dietary fibre intake has been associated with a reduced risk of obesity and metabolic disease such as type 2 diabetes [1]. A key mechanism that has been characterized in driving this protective effect is the enhanced secretion of the satiety-inducing gut hormones, Glucagon-like peptide 1 (GLP-1) and Peptide YY (PYY) following high fibre consumption [2–4].GLP-1 and PYY are released by enteroendocrine cells predominantly enriched in the distal gastrointestinal tract known as ‘L-cells’, in response to a range of nutrient and bacterial stimuli, including short chain fatty acids (SCFAs) [5,6]. SCFAs are metabolites derived from the anaerobic bacterial fermentation of dietary fibre by the gut microbiome and are both an energy source and ligands of G-protein coupled receptors (GPCRs) free fatty acid receptor 2 (FFAR2) and 3 (FFAR3) [7,8]. Acetate, propionate and butyrate are the predominant SCFAs in the human gut and activates human FFAR2 and FFAR3 with distinct potencies (human FFAR2; Ace = Pro > But, FFAR3; Pro = But > Ace) [7,8]. FFAR2 exhibits dual coupling capabilities to Gαi and Gαq heterotrimeric G protein pathways, whereas FFAR3 predominantly activates Gαi signalling [7,8]. While targeted colonic delivery of propionate-inulin ester has known beneficial actions in regulating appetite and weight gain in humans [9], FFAR2 has been demonstrated to be the primary receptor in driving GLP-1 and PYY release in response to SCFA activation in rodent models as of yet [10–12].

The mechanisms driving acute anorectic gut hormone release by propionate-activated FFAR2 have implicated both calcium and Gαi/p38 signalling in driving acute GLP-1 release from mouse L-cells [10,12]. However, SCFAs mediate multiple functions, beyond acute anorectic gut hormone release, not only across cell types but within L-cells [13–15]. How each SCFA elicits distinct physiological functions from FFAR2/3 that share common upstream G protein signal machinery is poorly understood [11]. One mechanism GPCRs employ is spatial pleiotropy in signalling, which for numerous GPCRs has been shown to be integral in achieving diversity and precision in their ligand-induced physiological effects [16]. We have previously demonstrated a key role for FFAR2 internalization and endosomal localization for propionate-mediated Gαi/p38 signalling in mouse L-cell models and colonic crypts [12].

In addition to hormone secretion, the response of L-cells to intestinal metabolites also influences their differentiation trajectories within the intestinal stem niche [17]. For example, L-cells derived from mice fed a high-fat diet have been shown to exhibit downregulated expression of nutrient-sensing machinery and enteroendocrine-specific transcription factors [18]. In contrast, supplementation with SCFAs has upregulated transcription factor cascades involved in L-cell development, and increased L-cell number in mouse and human organoids [19]. In human studies, this inherent ‘plasticity’ in L-cell differentiation has been demonstrated in obese individuals who exhibit a depleted density of GLP-1 and PYY-positive EECs that is restored upon laparoscopic sleeve gastrectomy [20,21]. EECs, like all intestinal cell types, are derived from the differentiation of multipotent stem cells. The primary checkpoint for a stem cell progenitor destined for an EEC fate is antagonized by the Notch effector protein Hes1 [22]. Hes1 has been found to be upregulated in the obese intestine and similarly reversed upon weight loss surgery [20] . Alongside local signalling gradients such as Wnt, Notch and BMP that co-ordinate the ISC niche, several GPCRs have been shown to regulate intestinal cell proliferation and differentiation [23]. SCFA-mediated activation of FFAR2 has been linked to increased numbers of PYY-positive L-cells by employing knockout mouse models [24], with indication that the SCFA butyrate may drive PYY transcription through poorly understood receptor and non-receptor mechanisms [25]. The signalling cascades and transcriptional function(s) of butyrate-mediated FFAR2 signalling in the human L-cell, remains unclear.

Here, we demonstrate that in a human L-cell model, SCFAs activate FFAR2 signalling pathways that vary highly with chain length of the fatty acid and exhibit differential location bias. This location bias in G protein signalling in turn mediates the ligand-specific actions of each SCFA on downstream L-cell functions, specifically at a transcriptional, morphological and endocrine level. Furthermore, we demonstrate that the SCFA butyrate promotes Notch-sensitive enteroendocrine differentiation to a ‘PYY-bias’ phenotype that can be augmented via a selective FFAR2 allosteric biased ligand, highlighting a critical role for butyrate in driving specific steps of L-cell differentiation via a spatially regulated FFAR2-G*α*i axis.

## Results

### The SCFA butyrate induces distinct endocrine and morphological changes in the human colonic enteroendocrine NCI-H716 cell line

The NCI-H716 cell line is a widely used model of human L-cells, particularly in studying nutrient-stimulated GLP-1 and PYY secretion [26,27]. We confirmed gene expression of both SCFA GPCRs *Ffar2* and *Ffar3* in this cell line, alongside GLP-1 precursor proglucagon (*Gcg)* and *Pyy* (**Figure S1).** In addition, previously reported [25] SCFA-mediated increases in gene expression of *Gcg* and *Pyy* were confirmed (**Figure 1A**). NCI-H716 cells were treated with 2 mM of acetate, propionate or butyrate for 24 h. Propionate and butyrate, but not acetate, significantly decreased *Gcg* expression. In contrast, all three SCFAs significantly increased *Pyy* transcript abundance, and this effect positively correlated with chain length (C2<C3<C4) with butyrate stimulating the largest increase in expression. This dramatic increase induced by butyrate on *Pyy* transcript was also reflected at the secretory level, however acetate and propionate did not increase PYY secretion in NCI-H716 cells (**Figure 1B**). Using immunofluorescent staining and confocal imaging paired with adaptive deconvolution, achieving a resolution limit of 120nm, GLP-1 and PYY granules were resolved to distinct populations in NCI-H716 cells (**Figure S1).** In untreated cells, PYY granules were less abundant than GLP- 1 granules, on average comprising 19.01 ± 2.138% of the total granule (GLP-1 and PYY) pool per cell (Figure 1C-D). Following butyrate treatment there was an increase in the total granule number (GLP-1 and PYY granules), and this was accompanied by a significant increase in %PYY granules to 34.26 ± 4.456 (**Figure 1E**).

**Figure 1:**
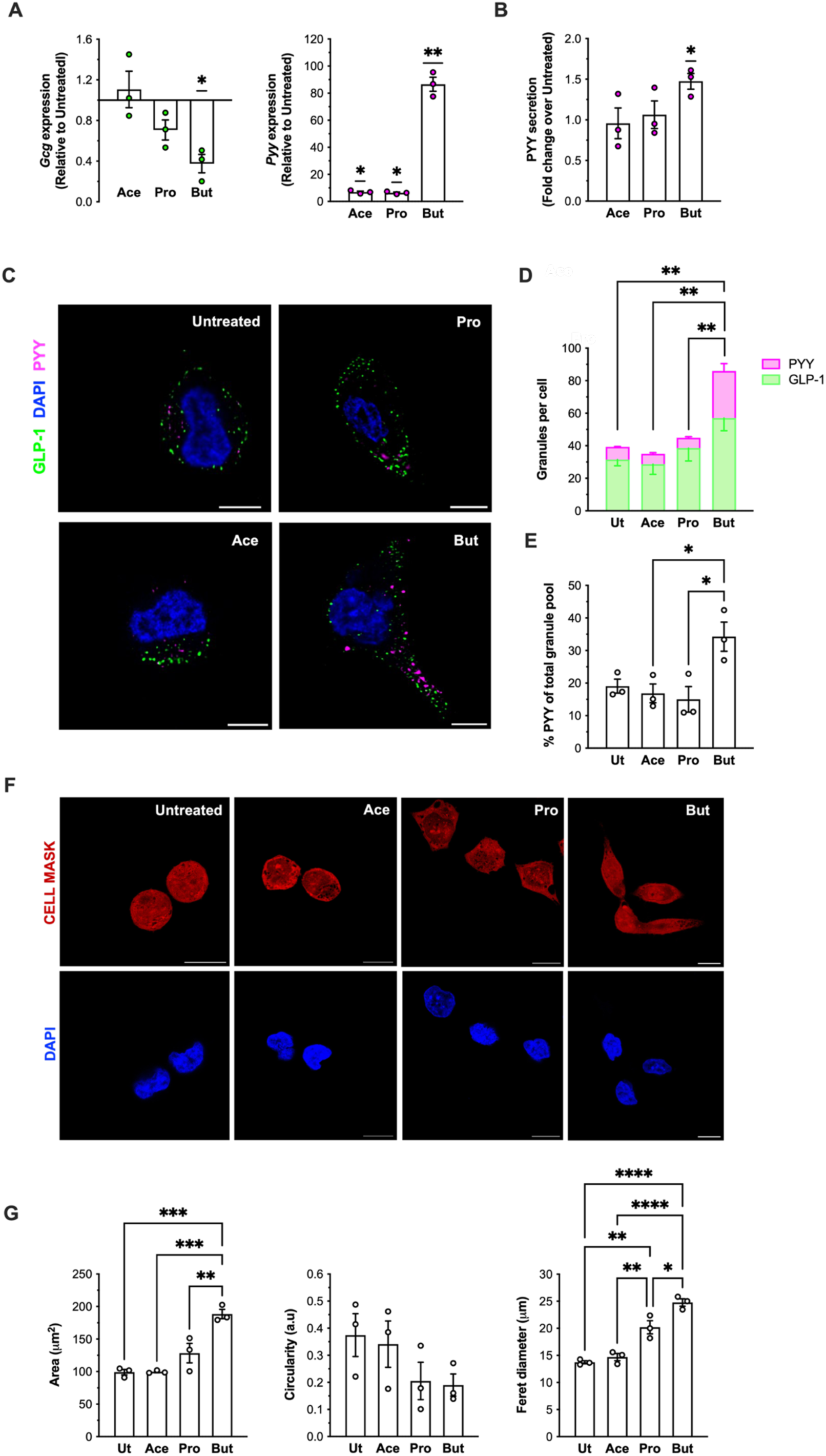
Butyrate induces enhances *Pyy* expression, PYY secretion and granule number, and induces morphological changes in NCI-H716 cells. NCI-H716 cell were treated with either 2 mM of sodium acetate (Ace), sodium propionate (Pro) or sodium butyrate (But) for 24 h. (A) *Gcg* and *Pyy* transcript expression in NCI-H716 cells were detected by RT-qPCR. Gene expression relative to untreated was determined using the 2^--ΔCt^ method with ribosomal protein L12 (*Rpl12)* as a housekeeping gene control. Data are presented as the mean ± SEM of data collected across three independent experiments in which conditions were run in at least triplicate. Symbols depict the mean ± SEM of individual biological repeats. (**p<0.01;*p<0.05, t-test SCFA vs Untreated). **(B)** Secretion of PYY from NCI-H716 cells was assayed by ELISA. Following SCFA treatment as for (A), cells were incubated with ligand-free secretion buffer for 2 h and supernatants were subsequently assayed for PYY concentration by ELISA. Data was normalised as fold change over untreated cells. Data are presented as the mean ± SEM of data collected across three independent experiments in which conditions were run in at least triplicate. Symbols depict the mean ± of individual biological repeats. (*p<0.05, t-test SCFA vs Untreated). **(C)** Representative images of fixed NCI-H716 stained with anti-GLP-1 and anti-PYY antibodies captured by confocal microscopy in super-resolution by an adaptive deconvolution module (LIGHTNING, Leica). Scale bar= 10μm. Inset scale bar= 2 μm. Conditions were imaged in at least triplicates across 5 independent experiments. **(D)** Total granule number, representing the sum of GLP-1 and PYY granules, in Untreated (Ut) or SCFA-treated NCI-H716 cells in super-resolved confocal images. Data represent the mean ± SEM of three independent experiments across which n=50 cells were analysed (**p<0.01; One-way ANOVA with Tukey’s multiple comparisons; Ut vs Ace vs Pro vs But). **(E)** % PYY granules of total (GLP-1 + PYY) granule pool Untreated (Ut) or SCFA-treated NCI-H716 cells in super-resolved confocal images. Data represent the mean ± SEM of three independent experiments across which n=50 cells were analysed (*p<0.05; One-way ANOVA with Tukey’s multiple comparisons; Ut vs Ace vs Pro vs But). **(F)** Representative images of fixed NCI-H716 cells captured via confocal microscopy incubated with HCS CellMask Deep Red Stain (stains across the cytoplasm and nuclei) and DAPI nuclei stain. Scale bar= 10μm. Conditions were imaged in at least quadruplicates across 3 independent experiments. **(G)** Measurements of mean area, circularity and feret diameter obtained by automatic particle analysis on thresholded images of Untreated (Ut) and SCFA-treated NCI-H716 cells by ImageJ. Data represent the mean ± SEM of three independent experiments across which n=120 cells were collectively analysed. Symbols depict the mean ± SEM of individual biological repeats. (****p<0.0001; ***p<0.001; **p<0.01; *p<0.05, One-way ANOVA with Tukey multiple comparisons; Ut vs Ace vs Pro vs But). See also Figure S1.

NCI-H716 cells treated with butyrate also displayed a prominent morphological shift; exhibiting elongated spindle-like morphology (**Figure 1F**). A CellMask stain was employed to quantify SCFA-mediated changes in morphological parameters such as cellular area, feret diameter (defined as the longest distance across the cell in any direction), and circularity (Perimeter^2^/4*π* x Area) (**Figure 1G**). We observed a significant increase in feret diameter amongst propionate- and butyrate-treated cells, with butyrate producing a significantly larger effect than propionate. Only butyrate significantly increased cellular area. Acetate had no effect on NCI-H716 morphology. Together these findings suggest that butyrate not only functionally drives L-cells towards production of PYY at a transcriptional, translational and secretory level, but also induces a morphology akin to L-cells in the colon [28,29].

### Butyrate activates distinct G-protein signalling profiles compared to acetate and propionate *via* FFAR2 in human L-cells

To uncover the mechanistic pathways underlying the reprogramming actions of butyrate in NCI- H716 cells, we characterized and compared the FFAR2/3 G protein signalling profiles following treatment with each SCFA. FFAR2 can signal via Gαi and Gαq pathways. Activation of Gαq stimulates the activity of phospholipase C-β, which cleaves PIP2 to produce DAG and IP3 leading to release of calcium from intracellular stores. We investigated SCFA-mediated activation of Gαq in the human NCI-H716 cells by measurement of IP1 levels, a downstream metabolite of IP3 (**Figure 2A**). We observed concentration-dependent activation of Gαq signalling following stimulation with either acetate (pEC50 = 3.403 ± 0.342) or propionate (pEC50 = 3.434 ± 0.225), which was inhibited by the FFAR2-selective orthosteric antagonist GLPG0974 [30] (**Figure 2B**). In comparison, butyrate stimulated a weaker increase in intracellular IP1 levels (pEC50 = 2.967 ± 0.219). Gαq activation profiles of SCFAs in NCI-H716 cells were also assessed by measurement of intracellular calcium mobilization (**Figure 2C, D**). Acetate and propionate triggered a rapid, monophasic increase in intracellular calcium. Butyrate failed to induce significant calcium mobilization, supporting minimal/weak activation of Gαq signalling in comparison to acetate and propionate.

**Figure 2:**
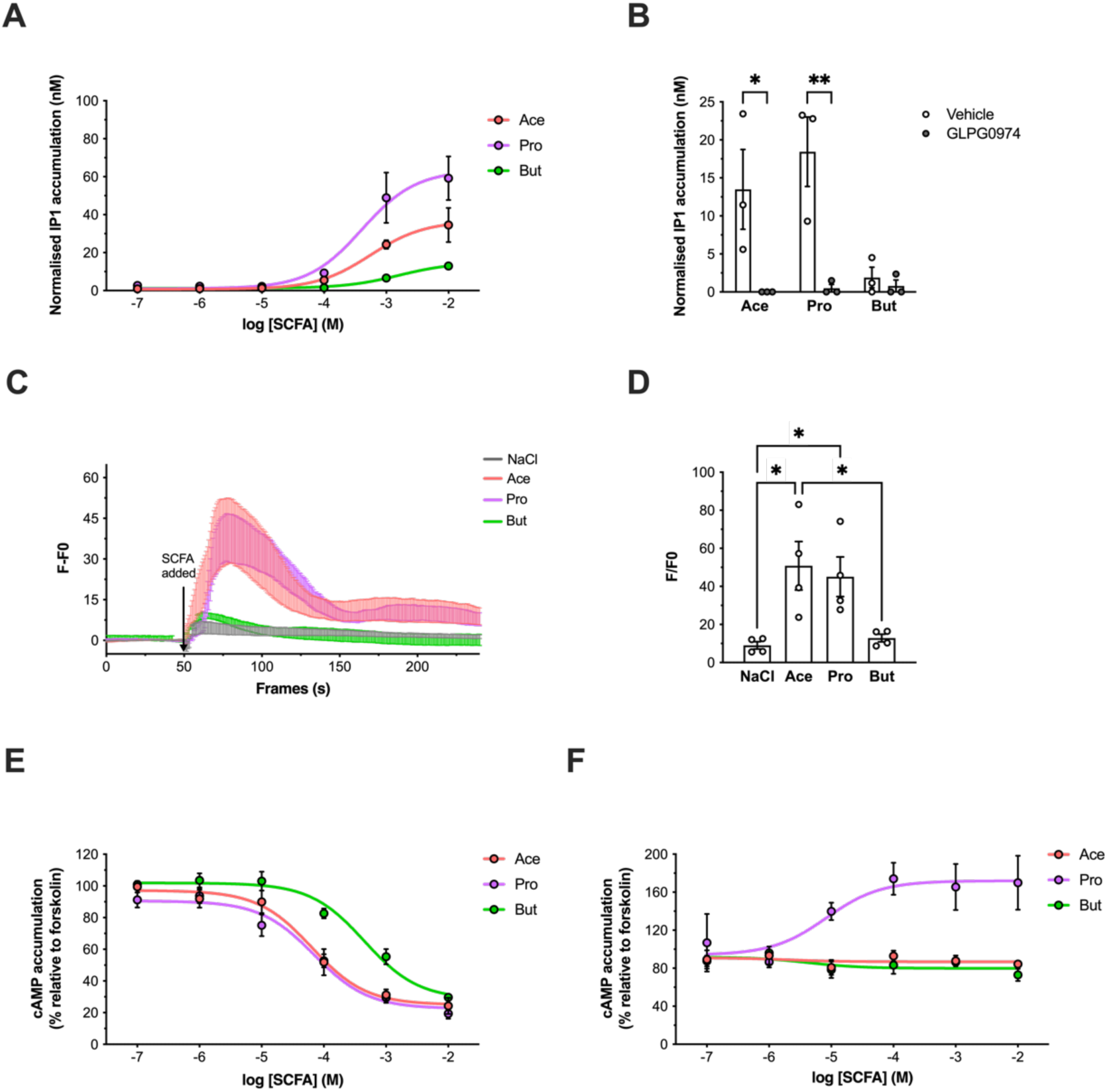
SCFAs stimulate distinct Gαi/o and Gαq/11 signalling profiles in NCI-H716 cells *via* activation of FFAR2. **(A)** Intracellular IP1 accumulation measured in NCI-H716 cells following 2h incubation with increasing concentrations of either NaCl, sodium acetate (Ace), sodium propionate (Pro) or sodium butyrate (But). Data were normalized by subtraction of NaCl response at each concentration. Data represent the mean ± SEM of three independent experiments in which conditions were run in at least triplicates. **(B)** IP1 accumulation in NCI-H716 cells pre-treated with DMSO (Vehicle) or FFAR2 antagonist GLPG0974 (1µM, 15 min) followed by 2h incubation with 1mM of NaCl, Ace, Pro or But. Data were normalized by an identical method to (A). Data are presented as the mean ± SEM of data collected across three independent experiments in which conditions were run in at least triplicates. Symbols depict the mean ± SEM of individual biological repeats. (**p<0.01;*p<0.05, t-test Vehicle vs GLPG0974 response). **(C & D)** Intracellular Ca^2+^ increases in NCI-H716 cells measured following 1h incubation with Fluo-4AM fluorescent dye, Cells were imaged live by confocal microscopy at 1.2 frames/second, at basal for 1 min, and then 10mM of NaCl, Ace, Pro or But was added. (C) Plot showing maximal fluorescent intensity following ligand addition normalised to untreated baseline on an individual cell basis (F-F0). Data are shown as the mean ± SEM of four independent experiments in which n=40 cells were analysed per experiment. Symbols depict the mean ± SEM of individual biological repeats. (*p<0.05, One-way ANOVA with post-hoc Tukey test; NaCl vs Ace vs Pro vs But). **(D)** Representative F-F0 traces of (C) from the beginning (0 s) to end (270 s) of experimental capture. **(E)** Inhibition of intracellular cAMP accumulation in NCI-H716 cells treated with 500nM 3-isobutyl-1- methylxanthine (IBMX) for 5 min and then stimulated with 3µM of forskolin (FSK) in the presence of increasing doses of Ace, Pro or But for 5 min. Data are expressed as % response of cAMP accumulation in FSK-treated cells and represent the mean ± SEM of five independent experiments in which conditions were run in triplicates. **(F)** Inhibition of cAMP accumulation in NCI-H716 cells following pre-treatment with DMSO (Vehicle) or GLPG0974 antagonist (1µM, 15 min) and subsequent ligand stimulation as in (E). Data represent the mean ± SEM of three independent experiments in which conditions were run in triplicates.

Activation of G⍺i signalling by GPCRs inhibits adenyl cyclase activity, thereby downregulating the conversion of ATP to cAMP and activity of cAMP-dependent protein kinase (PKA). NCI- H716 cells were treated with the adenyl cyclase activator forskolin, and the effect of SCFAs on forskolin-mediated cAMP accumulation was measured. All three SCFAs exhibited a significant inhibition of forskolin-mediated cAMP accumulation in a concentration-dependent manner, with butyrate (pIC50 = -3.360 ± 0.416) exhibiting significantly lower potency than propionate (pIC50 = -4.241 ± 0.523) and acetate (pIC50 = -4.230 ± 0.46) (**Figure 2E**). Given that both FFAR2 and FFAR3 are Gαi/o-coupled and both receptors are detected at mRNA level, it was important to resolve the mechanistic involvement of each receptor in driving G⍺i signalling. The selective FFAR2 antagonist GLPG0974 inhibited the G⍺i signalling across all doses of all three SCFAs employed, although propionate-induced an unexpected increase in cAMP signals following treatment with GLPG0974. Overall, these data suggest that FFAR2 rather than FFAR3 mediates SCFA-driven G⍺i responses in NCI-H716 cells **(Figure 2F**).

### Butyrate and a selective FFAR2 allosteric agonist increase *Pyy* secretion in an internalization-independent manner

We have previously demonstrated that in mouse enteroendocrine L-cells and colonic crypts, propionate-stimulated GLP-1 release is dependent on the internalization of FFAR2 for Gαi signalling [12]. An understanding of SCFA-encoded trafficking profiles of human FFAR2 is limited and could be crucial to unravelling functions specific to FFAR2 activation by butyrate in human L-cells. NCI-H716 cells were transfected with a FLAG-tagged human FFAR2 (hFFAR2) receptor, and receptor internalization was visualized by ‘feeding’ live cells with anti-FLAG antibody to specifically label cell surface receptor to track its endocytosis, followed by confocal microscopy imaging. Under basal conditions, receptor was primarily at the plasma membrane with some endosomes apparent indicating constitutive internalization, as observed for mouse FFAR2 [12]. However, treatment with acetate, propionate or butyrate resulted in an increase in the abundance of FFAR2 endosomes (**Figure 3A**). Both constitutive and propionate-induced internalization of mouse FFAR2 proceeds *via* a dynamin-dependent mechanism [12], in which the GTPase dynamin facilitates the intracellular scission of receptor-populated clathrin-coated pits at the plasma membrane. We confirmed both constitutive and SCFA-induced trafficking of hFFAR2 in NCI-H716 cells is also dynamin-dependent as pre-treatment with dynamin inhibitor Dyngo-4a inhibited receptor internalization (**Figure 3A**).

**Figure 3:**
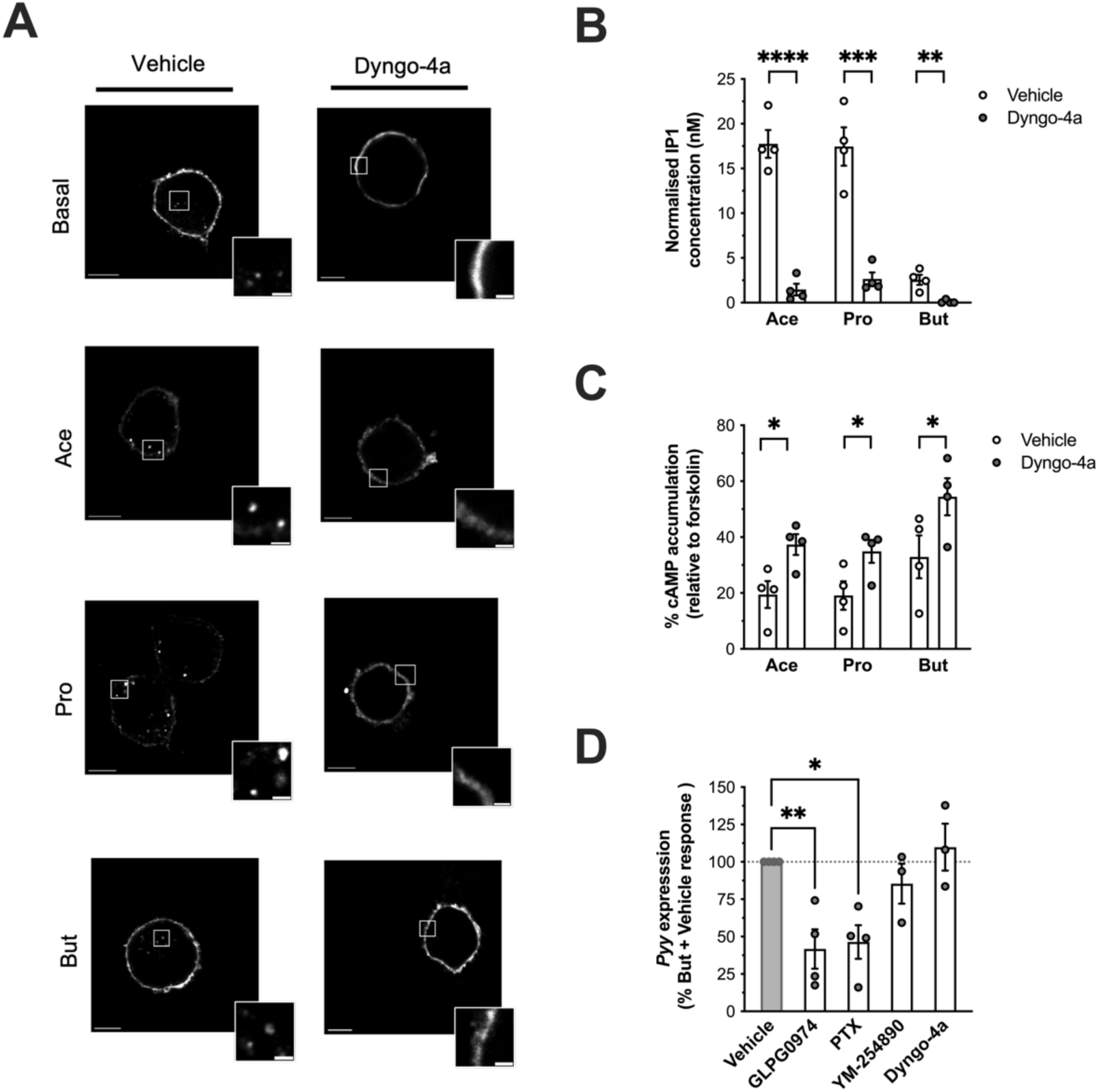
SCFAs stimulate dynamin-dependent internalization of FFAR2 from the plasma membrane, which modulate receptor signalling profiles. **(A)** Representative confocal images of NCI- H716 cells transiently transfected with FLAG-tagged human FFAR2. Transfected cells were treated for 45 min with DMSO (Vehicle) or 50 μm of dynamin-dependent endocytosis inhibitor Dyngo-4a, followed by staining with anti-FLAG M1 antibody for 30 min. 1mM of sodium acetate (Ace), sodium propionate (Pro) or sodium butyrate was added to cells in the latter 20 minutes of staining. Three independent experiments were carried out, and n=10 cells were captured per experiment. Scale bar= 10μm. Inset scale bar= 2 μm. **(B)** Intracellular IP1 accumulation in NCI-H716 cells measured following 45 min pre-treatment with DMSO (Vehicle) or 50μM of Dyngo-4a and then a 2hr stimulation with 1mM of NaCl, Ace, Pro or But. Data were normalized by subtraction of NaCl response at each concentration. Data represent the mean ± SEM of four independent experiments in which conditions were run in at least triplicates. Symbols depict the mean ± SEM of individual biological repeats. (****p<0.0001, ***p<0.001, **p<0.01, t-test Vehicle vs Dyngo-4a response). **(C)** Intracellular cAMP accumulation measured in NCI-H716 cells pre-treated for 45 min with DMSO (Vehicle) or 50μM Dyngo-4a and then stimulated for 5 min with 500nM 3-isobutyl-1- methylxanthine (IBMX) followed by 3μM of forskolin (FSK) in the presence of 1mM of Ace, Pro or But for 5 min. Data are expressed as % response of cAMP accumulation in FSK-treated cells and represent the mean ± SEM of four independent experiments in which conditions were run in triplicates. Symbols depict the mean ± SEM of individual biological repeats. (*p<0.05, t-test Vehicle vs Dyngo-4a response. **(D)** Butyrate-mediated upregulation of *Pyy* transcript expression measured in NCI-H716 cells pre-treated with DMSO (Vehicle), GLPG0974 (1μM, 15 min), PTX (500ng/ml, 20 hours), Gαq inhibitor YM-254890 (10nM, 15 min) or dynamin-dependent endocytosis inhibitor Dyngo-4a (50uM, 45 min) followed by 24hr incubation with 2mM of But and detected by RT-qPCR. Relative gene expression was determined as in (A). But + Inhibitor response is shown as % percentage of But + Vehicle response (dotted line at 100%). Data are presented as the mean ± SEM of data collected across three independent experiments in which conditions were run in at least triplicate. Symbols depict the mean ± SEM of individual biological repeats. (**p<0.01, *p<0.05; One-way ANOVA with Dunnett’s post-hoc test, But + Vehicle (control) vs But + inhibitor). See also Figure S2.

To uncover the role of SCFA-induced hFFAR2 internalization on receptor signalling profiles in NCI-H716 cells, Gαq and Gαi signalling was measured following treatment with Dyngo-4a. A strong inhibition of both acetate- and propionate-stimulated increases in IP1 accumulation by Dyngo-4a was observed, indicating that internalization of hFFAR2 was required for full activation of Gαq (**Figure 3B**). Although butyrate’s ability to activate Gαq is limited, the small increase in IP1 levels was also inhibited by Dyngo-4a. SCFA-mediated activation of Gαi signalling was only partially inhibited by Dyngo-4a (**Figure 3C**). These findings demonstrate that hFFAR2 exhibits a differential requirement for internalization for signalling via each G protein pathway by SCFAs.

As butyrate activates primarily Gαi signalling via an FFAR2-dependent mechanism in NCI-H716 cells, potentially from both the plasma membrane and following receptor internalization, we next assessed the mechanistic involvement of this pathway in driving expression of *Pyy*. Following pre-treatment with the FFAR2 antagonist GLPG0974, butyrate-mediated *Pyy* upregulation was partially but significantly inhibited (**Figure 3C**). Likewise, pre-treatment with the Gαi inhibitor pertussis toxin (PTX), at a concentration which we had observed to significantly diminish Gαi responses activated by up to 10mM of butyrate **(Figure S2),** partly but significantly inhibited butyrate-mediated Pyy gene upregulation. In contrast, treatment with an inhibitor of Gαq, YM- 275890, did not affect butyrate-mediated increases in *Pyy* transcript, supporting the involvement of an FFAR2-Gαi axis in this downstream response. Although FFAR2-Gαi signalling in NCI- H716 cells was partially dependent on receptor internalization, Dyngo-4a treatment did not inhibit butyrate-mediated *Pyy* upregulation, indicating a plasma membrane hFFAR2-Gαi signal in driving this response. In contrast to our findings on *Pyy* expression, butyrate-stimulated PYY secretion was not inhibited by GLPG0974 and signalling inhibitors, indicating a potential FFAR2- independent mechanism. **(Figure S2).**

To further investigate a role for an FFAR2-Gαi pathway in driving *Pyy* expression, we employed a selective hFFAR2 allosteric biased agonist of Gαi activation, AZ-1729 [31]. Stimulation of NCI-H716 cells with AZ-1729, with or without butyrate, resulted in activation of Gαi signalling and potentiation of butyrate responses, confirming its ability as an agonist and positive allosteric modulator **(Figure S3).** In NCI-H716 cells expressing FLAG-hFFAR2, we observed dynamin-dependent receptor internalisation following treatment with AZ-1729, which was blocked by Dyngo-4a pre-treatment (Figure 5A). Dyngo-4a did not, however, significantly impact AZ-1729- mediated Gαi signalling in NCI-H716 cells, suggesting that the FFAR2-Gαi signal pathway activated by AZ-1729 is internalization-independent and occurs at the plasma membrane (**Figure 4B**). AZ-1729 stimulated a ∼2-fold increase in *Pyy* expression relative to baseline, however when co-treated with butyrate increased *Pyy* mRNA levels a further ∼2 fold over the ∼16 fold response driven by butyrate, suggesting a marked synergistic action (**Figure 4C-D**). Together, these findings not only reinforce the mechanistic involvement of a plasma membrane-localised hFFAR2-Gαi signalling pathway in *Pyy* expression but that an allosteric modulator can amplify the actions of butyrate in L-cells.

**Figure 4:**
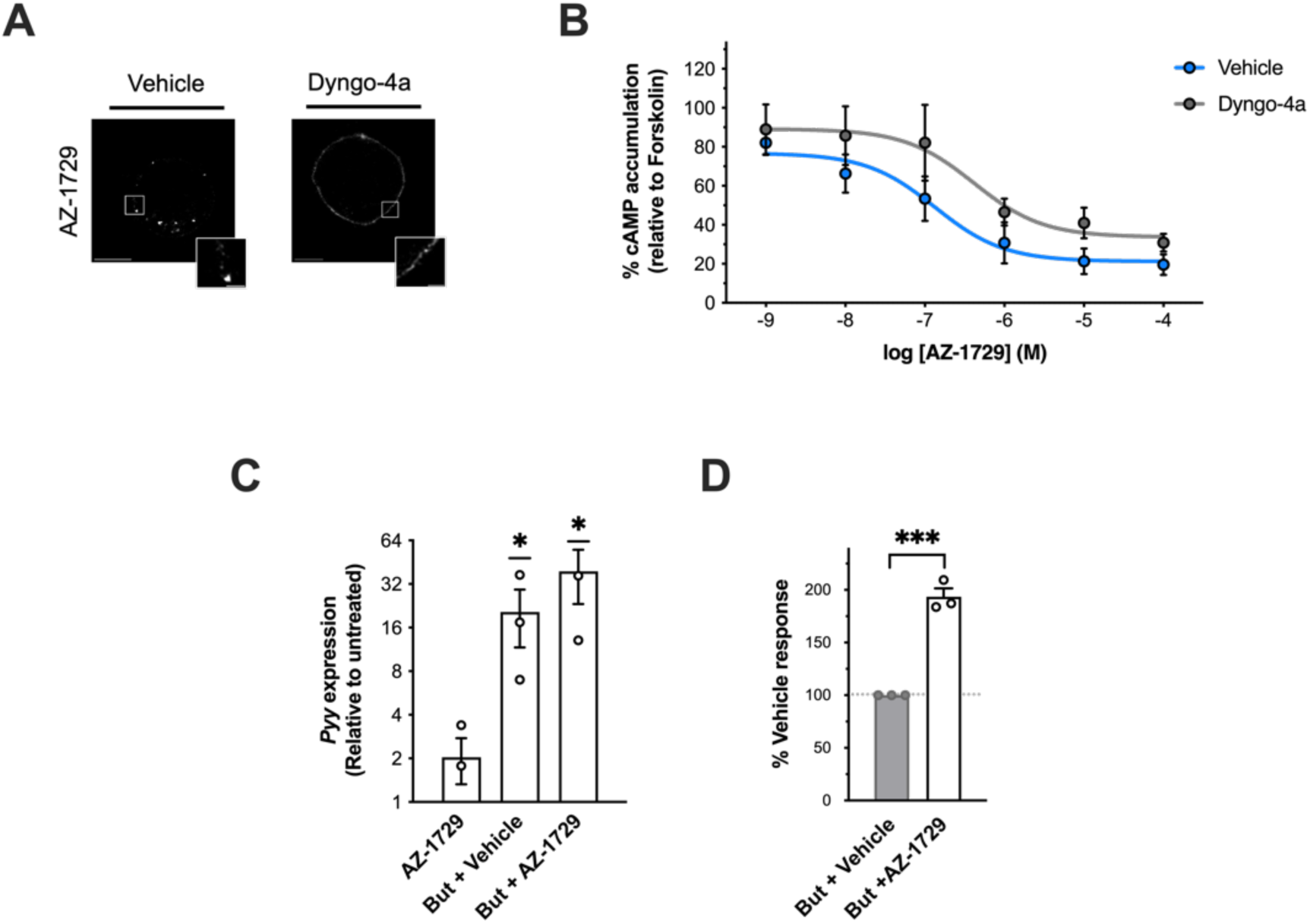
AZ-1729 activates FFAR2-Gαi signalling independent of receptor internalization to upregulate *Pyy* expression. **(A)** Representative confocal images of NCI-H716 cells transiently transfected with FLAG-tagged human FFAR2. Transfected cells were treated for 45 min with DMSO (Vehicle) or 50μM of dynamin-dependent endocytosis inhibitor Dyngo-4a, followed by staining with anti-FLAG M1 antibody for 30 min. FFAR2-Gαi biased allosteric agonist AZ-1729 (1μM, 20 min) was added to cells. Three independent experiments were carried out, and n=5 cells were captured per experiment. Scale bar= 10μm. Inset scale bar= 2 μm. **(B)** Intracellular cAMP accumulation measured in NCI-H716 cells pre-treated for 45 min with DMSO (Vehicle) or 50 μM Dyngo-4a and then stimulated for 5 min with 500 nM 3-isobutyl-1- methylxanthine (IBMX) followed by 3 μM of forskolin (FSK) in the presence of 1 μM of AZ-1729 for 5 min. Data are expressed as % response of cAMP accumulation in FSK-treated cells and represent the mean ± SEM of three independent experiments in which conditions were run in triplicates. **(C & D)** *Pyy* transcript expression in NCI-H716 cells treated for 24h with 1 μM AZ-1729, 2mM sodium butyrate (But) + DMSO (Vehicle) or a combination of 2 mM But and 1 μM AZ-1729. Transcript levels were detected by RT-qPCR and gene expression relative to untreated was determined using the 2^--ΔCt^ method with ribosomal protein L12 (*Rpl12)* as a housekeeping gene control. In (D), But + AZ-1729 response is shown as % of But + Vehicle response (dotted line at 100%). Data are presented as the mean ± SEM of data collected across three independent experiments in which conditions were run in at least triplicate. Symbols depict the mean ± SEM of individual biological repeats. (*p<0.05, One-sample Control value (1) vs Ligand(s); ***p<0.01, t-test But + Vehicle vs But + AZ-1729). See also Figure S3.

**Figure 5:**
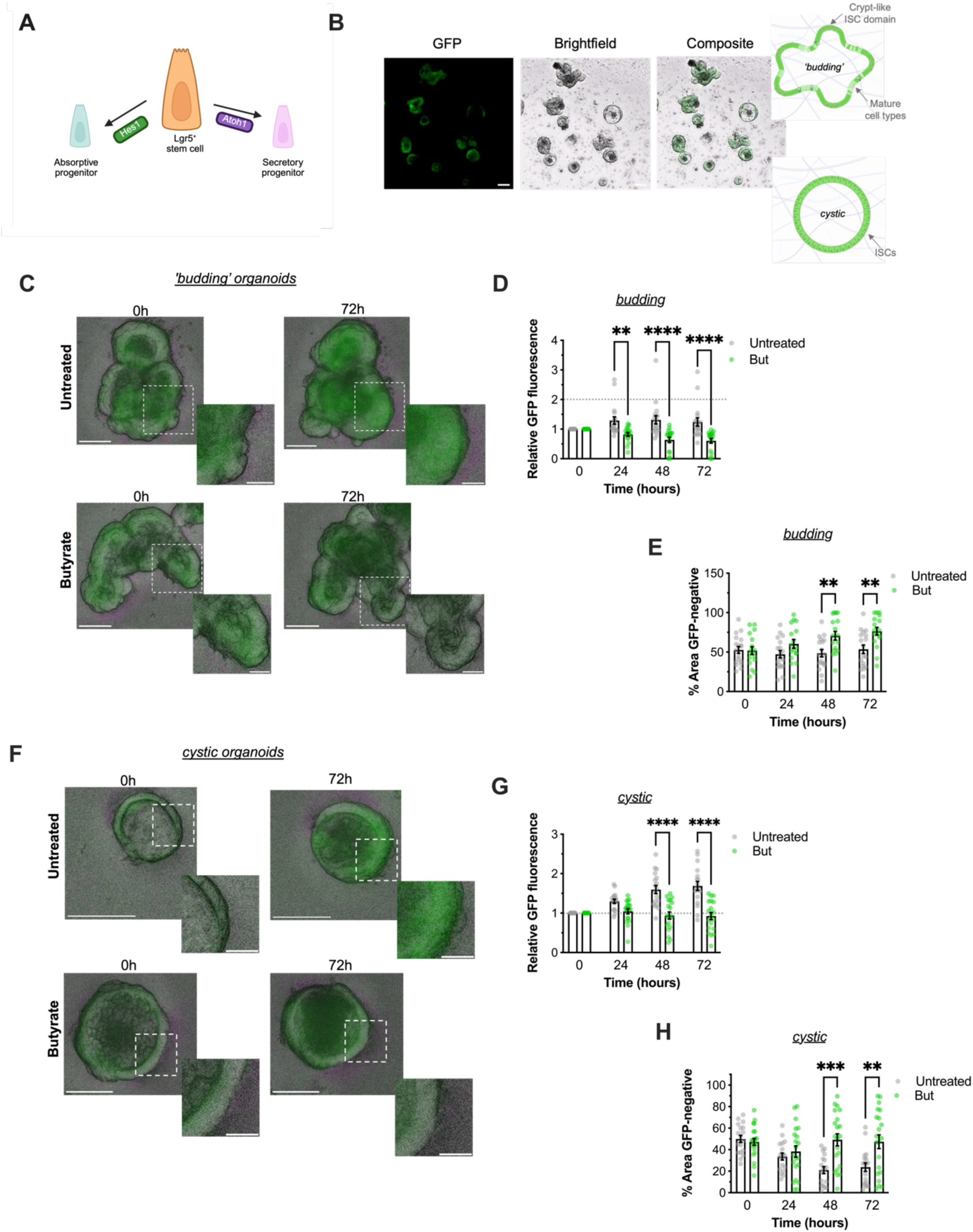
Butyrate dampens Notch activity in Hes1-GFP mouse colonic organoids. **(A)** Cartoon depicting the Notch-modulated cell fate checkpoint involved in Lgr5+ stem cell differentiation to an absorptive or secretory fate *via* Hes1 or Atoh1 respectively. **(B)** Representative images of colonic Hes1- GFP mouse organoids in culture captured live by a widefield fluorescent microscope. Cartoons of the cellular architecture of ‘budding’ organoids, consisting of mature cell types and crypt-like domains populated by intestinal stem cells (ISCs) and mature cell types, and cystic organoids rich in ISCs. **(C-H)** Hes1-GFP organoids were monitored across a 72h timeframe in Untreated culture conditions or cultures incubated with 2 mM of sodium butyrate (But) for 24 h. Analyses of ‘budding’ and ‘cystic’ organoids were carried out separately. Data from n=18 ‘budding’ organoids and n=19 ‘cystic’ organoids were collected across cultures from three separate Hes1-GFP mice. **(C)** Representative images of ‘budding’ organoids. Scale bar= 60μm. Inset scale bar= 25μm. **(D)** Relative GFP fluorescence of individual ‘budding’ organoids shown as fold change of fluorescence at 0h at each time point. **(E)** GFP-negative area of individual ‘budding’ organoids shown as % of total organoid area. **(F)** Representative images of cystic organoids. Scale bar= 60μm. Inset scale bar= 15μm**. (G)** Relative GFP fluorescence and **(H)** % GFP-negative area measurements of cystic organoids respectively. Symbols represent individual organoids. (****p<0.01, **p<0.01, *p<0.05; Two-way ANOVA with post-hoc Sidak’s test Untreated vs Butyrate). See also Figure S4.

### Butyrate modulates NOTCH activity to direct L-cell maturation

In addition to modulating the endocrine functionalities of L-cells in the intestinal milieu, SCFAs have been shown to influence EEC differentiation [19,24]. We next determined if the transcriptional, endocrine and morphological actions of butyrate reflect activation of key differentiation pathways driving L-cell development. In the intestinal stem cell niche (ISC), secretory lineage differentiation is antagonized by Notch. The Notch effector protein Hes1 is expressed in different cellular populations in the ISC niche where it serves distinct functions; in ISCs its promotes their continuous proliferation, whereas in progenitors cells it encourages differentiation to an absorptive (i.e., enterocyte) rather than a secretory fate (i.e., goblet, Paneth, EEC) via repression of the master secretory lineage development transcription factor, Atoh1 (**Figure 4A**) [32–34]. To investigate the role of butyrate on secretory lineage differentiation, we established colonic organoids from a previously described Notch reporter Hes1-GFP mouse [33].

Organoid cultures were established from isolated Hes1-GFP colonic crypts and comprised of a mixture of two canonical organoid populations with heterogeneous morphologies and cellular makeup; a ‘budding’ population and a ‘cystic’ population (**Figure 5B**). Mature “budding” organoids consist of projections of crypt-base-like domains, enriched in proliferative stem cells, and upper-crypt-like domains in which late-stage progenitors and mature differentiated cells of both absorptive and secretory lineages reside. As such, budding organoids recapitulate the *in vivo* architecture and functional compartmentalization of stem cell and differentiated zones in the intestinal colonic crypt [35–37]. In contrast, “cystic” organoids consist of a lumen surrounded by a thin epithelial monolayer of stem cells akin to populations of continuously dividing ISCs that occupy the base of crypts and support homeostatic renewal in the intestinal stem cell niche [38]. To understand whether differences in NOTCH activity (i.e., Hes1-GFP fluorescence) would be representative of heterogeneous cell ontologies in this reporter model, we analysed *Hes1* expression at single cell resolution in the mouse colon using the publicly available database Tabula Muris [39] (**Figure S4).** We confirmed *Hes1* expression was high across epithelial and enterocyte populations, and low in EEC and brush cells suggesting that differences in absorptive and secretory lineages could be delineated by reporter fluorescence.

We monitored the effect of butyrate on Hes1-GFP fluorescence in budding (**Figure 5C-E**) and cystic (**Figure 5F-H**) organoids across a 72h period. In budding populations, butyrate stimulated a significant time-dependent decrease in normalized fluorescence, which was not observed across untreated budding organoids (**Figure 5D**). We did not detect time-dependent differences in total area of budding organoids in either butyrate or untreated populations across timepoints **(Figure S4)**, however, areas devoid of GFP signal significantly increased in a time-dependent manner in butyrate-treated budding organoids (**Figure 5E**). In cystic organoids, GFP fluorescence significantly increased in a time-dependent manner in untreated conditions. Such an increase was not detected in butyrate-treated populations (**Figure 5G**). In addition, the percentage area that was GFP negative increased in a time-dependent manner in untreated cystic organoids, although this trend was also not detected in butyrate-treated equivalents (**Figure 5H**). Total area of cystic organoids was unchanged across time points in untreated conditions, yet was significantly increased following 72 h of butyrate treatment (**Figure S4).** Together, these observations in cystic and budding organoids illustrate an inhibitory effect of butyrate on Notch activity across ISCs and maturing cell types in the mouse colonic epithelium respectively.

### Butyrate-activated FFAR2 selectively modulates late stage maturation of EECs by increasing Pax6, a pathway amplified by AZ-1729

We then examined whether butyrate-mediated inhibition of Notch was indicative of enhanced cellular differentiation to secretory cell fates, and particularly to EECs. In EEC progenitors, *Atoh1* expression marks the initiation of a conserved cascade of transcription factors which are transiently and sequentially expressed, consisting of *Atoh1*, *Neurog3* and then *NeuroD1* [40–43] (**Figure 6A**). In NCI-H716 cells, incubation with butyrate significantly downregulated *Atoh1,* whereas limited effect on *Neurog3* transcript levels were detected (**Figure 6B**). Notably, butyrate significantly increased *NeuroD1* expression by ∼4 fold, however, a significant increase in *NeuroD1* was observed. When we compared the effects of butyrate on modulating expression levels of *Atoh1, Neurog3* and *NeuroD1* with acetate and propionate, we only detected a significant decrease in *Atoh1* expression by propionate (**Figure S5).**

**Figure 6:**
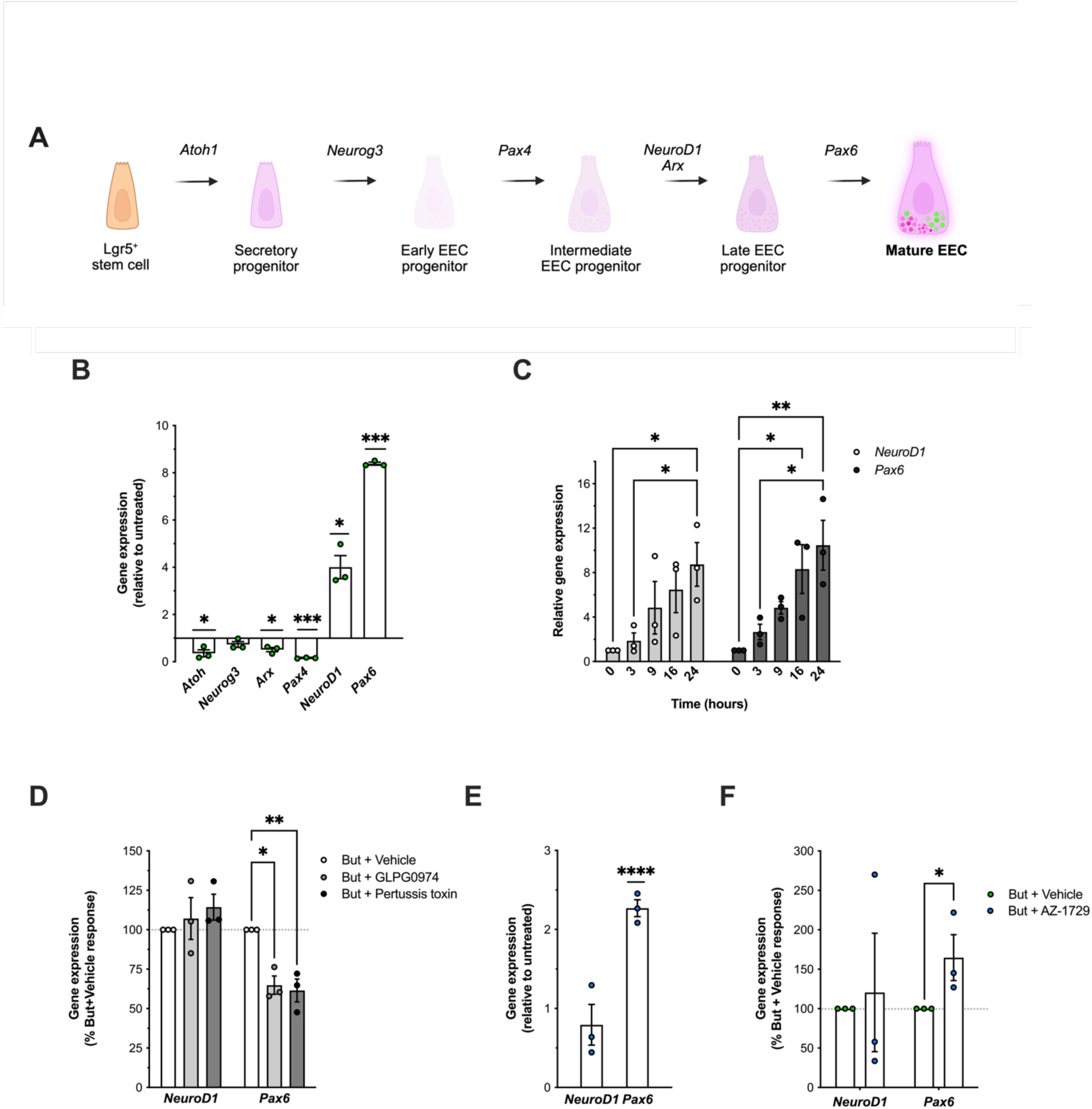
Butyrate and AZ-1729 modulate the expression of EEC-maturation transcription factors. **(A)** Cartoon depicting the signature transcription factor cascades involved in the differentiation of Lgr5+ stem cells to mature L-cells. **(B)** Relative gene expression of transcription factors involved in L-cell differentiation (*Atoh1, Neurog3, Arx, Pax4, NeuroD1* and *Pax6)* in NCI-H716 cells treated for 24h with 2mM of sodium butyrate (But). Transcript levels were detected by RT-qPCR and gene expression relative to untreated was determined using the 2^--ΔCt^ method with ribosomal protein L12 (*Rpl12)* as a housekeeping gene control. Data are presented as the mean ± SEM of data collected across three independent experiments in which conditions were run in at least triplicate. Symbols depict the mean ± SEM of individual biological repeats. (****p<0.0001, *p<0.05, One sample t-test; Untreated (1) vs Butyrate). **(C)** *NeuroD1* and *Pax6* transcript expression in NCI-H716 cells treated with 2mM of But for 3, 9, 16 or 24h. Transcript levels were detected by RT-qPCR and gene expression was determined as in (B) Data are presented as the mean ± SEM of data collected across three independent experiments in which conditions were run in at least triplicate. (**p<0.01, *p<0.05, Two-way ANOVA with post-hoc Tukey’s test, 0h vs 3h vs 9h vs 16h vs 24h). **(D)** *NeuroD1* and *Pax6* transcript expression in NCI-H716 cells pre-treated with DMSO (Vehicle), FFAR2 antagonist GLPG0974 (1uM, 15 min) or Gαi inhibitor pertussis toxin (PTX) (500ng/ml) followed by 24h incubation with 2mM of But. Transcript levels were detected by RT-qPCR and gene expression was determined as in (B). Data is shown as But + Inhibitor response as % of But + Vehicle response (dotted line at 100%) for individual gene targets. Data are presented as the mean ± SEM of data collected in three independent experiments in which conditions were run in at least triplicate. Symbols depict the mean ± SEM of individual biological repeats. (**p<0.001, *p<0.05, Two-way ANOVA with post-hoc Tukey’s test; Vehicle vs GLPG0974 vs Pertussis toxin). **(E)** NeuroD1 and Pax6 transcript expression in NCI-H716 cells treated for 24h with 1μM of AZ-1729 and detected by RT-qPCR. Relative gene expression was determined as in (B). Data are presented as the mean ± SEM of data collected across three independent experiments in which conditions were run in at least triplicate. Symbols depict the mean ± SEM of individual biological repeats. (****p<0.0001, One-sample t-test DMSO Vehicle (1) vs AZ-1729). **(F)** *NeuroD1* and *Pax6* transcript expression in NCI-H716 cells treated for 24h with 2mM But + DMSO (Vehicle) or a combination of 2mM But and 1μM AZ-1729 detected by RT-qPCR. Relative gene expression was determined as in (B), (C) and (D). But + AZ-1729 response is shown as % of But + Vehicle response (dotted line at 100%) for individual gene targets. Data are presented as the mean ± SEM of data collected across three independent experiments in which conditions were run in at least triplicate. Symbols depict the mean ± SEM of individual biological repeats. (*p<0.05, t-test But + Vehicle vs But + AZ-1729). See also Figure S6.

Downstream of the *Atoh1-Neurog3-NeuroD1* cascade, other transcription factors participate in specifying the differentiation into EEC sub-lineages. The activity of the homeodomain transcription factors *Pax4*, *Arx* and *Pax6* have specifically been mapped to the production of GLP+ and/or PYY+ EECs *in vivo* and *in vitro* [24,43–45]. Butyrate treatment led to a significant downregulation in *Arx* expression, with an even greater decrease observed for *Pax4* expression. In contrast, expression of the transcription factor *Pax6*, which has previously been characterized as a direct downstream target *NeuroD1*, was significantly upregulated by butyrate [46]. We measured a time-dependency in butyrate-mediated upregulation of both *NeuroD1* and *Pax6* transcript levels (**Figure 6C**). No significant effects of acetate or propionate on the *Atoh1- Neurog3-NeuroD1* cascade apart from a significant downregulation in *Pax6* expression by acetate (**Figure S5).** Together, findings demonstrate selective effects of butyrate on L-cell developmental cascades that are specific to later stages of differentiation.

As we had identified a mechanistic involvement of FFAR2-Gαi activation in butyrate-mediated upregulation of PYY secretion, we next investigated the role of this pathway in modulating transcriptional programs that regulate L-cell differentiation (**Figure 6D**). Neither GLPG0974 nor pertussis toxin inhibited butyrate-mediated upregulation of *NeuroD1* expression. In contrast, butyrate-mediated upregulation of *Pax6* expression was significantly depleted by both inhibitors. When we assessed the ability of AZ-1729 to modulate expression of both these targets, we observed a significant upregulation of *Pax6* but not *NeuroD1* further supporting a selective role of FFAR2-Gαi in *Pax6* expression (**Figure 6E**). Finally, we observed enhanced upregulation of *Pax6* only upon dual treatment with butyrate and AZ-1729 (**Figure 6F**). Together, these findings collectively support FFAR2-independent mechanisms function in regulating *NeuroD1* while the activation of an FFAR2-Gαi pathway by butyrate drives the upregulation of *Pax6* expression, and can be further enhanced pharmacologically with AZ-1729.

## Discussion

Enteroendocrine L-cells play key roles in regulation of appetite through production and secretion of anorectic gut hormones GLP-1 and PYY in response to nutrients and metabolites. In addition, the gut luminal environment is known to influence the plasticity of the gut epithelia, including the programming of enteroendocrine cell abundance and type. The SCFAs are known to drive both acute gut hormone release and selectively enhance levels of PYY positive L-cells at a transcriptional and translational level [24,25] via receptor dependent and independent pathways. However, little is known about how SCFA GPCRs selectively modulate transcriptional and endocrine profiles of L-cells. In this study, we identify that butyrate promotes maturation and lineage specification of L-cells to mature PYY positive L cells at a transcriptional, endocrine and morphological level. In human L-cell models we uncover a spatially controlled FFAR2–Gαi signalling mechanism that can be pharmacologically amplified to promote Pyy and L-cell maturation.

The SCFAs, and in particular butyrate, are known to strongly induce a PYY-enriched hormone repertoire in human and mouse L-cells, including the NCI H716 cells employed in this study. Furthermore, our data reveal a partial dependency (∼50%) on FFAR2 for butyrate-induced upregulation of *Pyy* expression, consistent with the magnitude of inhibition observed in FFAR2 knockout models previously reported [25], suggesting butyrate may exert some transcriptional regulation in intestinal epithelial cells *via* its functionality as a HDAC inhibitor [47,48]. However, using FFAR2-selective antagonists and allosteric biased agonists with profiling of the upstream G protein signal pathways activated by SCFAs in human L-cells, we propose a key role for a butyrate-mediated FFAR2–Gαi signalling axis in L-cell function. Moreover, we also demonstrate a concomitant transcriptional modulation of Notch-sensitive cascades involved in L-cell maturation by butyrate, and a specific dependency on FFAR2-Gαi signalling in upregulation of the late-stage transcription factor *Pax6*.

Although SCFAs activate both FFAR2 and FFAR3 and both receptors were detected in NCI- H716 cells at transcript level, our data demonstrate that FFAR2 mediates Gαi- and Gαq-coupled signalling responses to acetate, propionate, and butyrate in a human L-cell line. Consistent with findings from recombinant receptor overexpression systems [7,8], we observe a striking bias of butyrate toward Gαi activation, in contrast to acetate and propionate, which robustly engage both Gαi and Gαq pathways. This signalling preference may underlie the prominent FFAR2- dependent effects of butyrate on *Pyy* in human L-cells. Supporting this notion, the selective FFAR2-Gαi biased agonist AZ1729 also increased *Pyy* transcript levels, reinforcing the role of Gαi-biased signalling in this context. Notably, our findings propose a plasma membrane-localized mechanism of a FFAR2-Gαi axis in the regulation of *Pyy* expression by butyrate and AZ1729. In contrast, our previous work identified an intracellular mode of propionate-mediated FFAR2–Gαi signalling from an endosomal compartment, which mediated acute GLP-1 secretion in mouse L-cells [11].These findings raise the possibility that distinct SCFAs may exhibit location bias in FFAR2-Gαi signalling to diversify its actions from a common G protein signal pathway. Additionally, species-specific differences in receptor signalling may contribute to functional divergence in some L-cell responses, as evidenced by the distinct signalling and functional profiles of mouse and human FFAR2 in monocytes [49], although butyrate’s ability to modulate differentiation in both mouse and human intestinal in this study [40] may indicate that certain pathways/functions may be conserved.

We have previously demonstrated that inulin, a fermentable substrate for SCFA production by the gut microbiome, acts *via* FFAR2 to selectively promote the expansion of PYY-positive, but not GLP-1-positive, L-cell populations in mice [24]. Building on these findings, our current study reveals a direct role of butyrate in influencing L-cell differentiation, selectively promoting a PYY- enriched transcriptional and functional profile. The selective effect(s) of butyrate on PYY agree with previous reports of mutually exclusive secretion of GLP-1 or PYY from L-cells upon acute nutrient challenge [28,50–53], and might reflect induction of ’switching’ of hormonal repertoires of EEC lineages as they mature [43,54]. Beyond its similar anorectic actions to GLP-1, PYY modulates other gastrointestinal functions including the efficiency of digestion and nutrient absorption [55], and colonic epithelial proliferation [56,57]. High intakes of fermentable fibre are related to expansion of butyrogenic bacterial populations [58] and increased butyrate flux in the colon [59,60]. Selectively targeting PYY release from colonic L- cells, which have been primed by a butyrate dominant environment, might promote these benefits and enhance the positive effects of high fibre consumption, whilst simultaneously reducing the incidence of adverse effects that might occur with co-release of GLP-1 and PYY due to overstimulation of central and peripheral pathways. Notably, we observed a morphological shift in NCI-H716 cells involving the formation of long cytoplasmic processes resembling those of PYY-positive EECs previously reported within colonic crypts [28,29]. Prior studies have demonstrated that the basal processes of PYY-expressing enteroendocrine cells (EECs), now termed neuropods, in fact resemble axonal structures both anatomically and functionally; neuropods form synapse-like connections with neurons and the intestinal milieu, enabling direct communication between the gut epithelium and the nervous system [43–47]. The existence of PYY-positive neuropods in the human colonic epithelium are yet to be characterized and might represent a unique L-cell population that is maintained by luminal and/or circulating butyrate.

SCFAs have previously been shown to transcriptionally modulate Notch-sensitive cascades of L-cell differentiation in human organoids [19]. Importantly, Notch signalling in the intestinal epithelium promotes ISC proliferation and antagonizes their differentiation to secretory fates via its effector Hes1 [22,33]. Using a Hes1-GFP organoid line, we tracked the effects of butyrate on Notch-mediated proliferation and inhibition of secretory lineage commitment over time at single-organoid resolution. In ISC-rich cystic organoids, butyrate induced a sustained plateau in Notch activity compared to untreated controls. In contrast, more differentiated, budding organoids displayed a progressive decline in Notch activity over time following butyrate exposure. These dynamic patterns suggest that butyrate modulates enteroendocrine cell (EEC) differentiation in the intestinal stem cell (ISC) niche through two distinct mechanisms: first, by promoting the loss of stemness and initiating differentiation in immature ISCs, and second, by constraining the differentiation potential of post-mitotic progenitors and/or driving transdifferentiation of mature epithelial cells toward ‘Notch-low’ secretory fates.

Butyrate also modulates transcriptional programs in the human enteroendocrine cell line NCI- H716. Consistent with our organoid data, butyrate robustly upregulated expression of transcription factors (*NeuroD1* and *Pax6)* involved in the late stages of L-cell differentiation cascades that are typically suppressed by Notch signalling. Among the three major SCFAs, butyrate elicited the most pronounced transcriptional response. Mechanistically, we identified a selective role for FFAR2-Gαi signalling in mediating butyrate-induced upregulation of the L-cell-specific factor *Pax6.* In contrast, the induction of the pan-EEC transcription factor *NeuroD1* by butyrate occurred independently of FFAR2, implicating alternative pathways such as HDAC inhibition or activation of another candidate GPCR such as GPR109A/HCA2 [66]. Together, these findings underscore the complexity of butyrate’s action in fine-tuning EEC differentiation, revealing a dual mechanism through which it promotes PYY-biased L-cell fates via both FFAR2- dependent and -independent pathways. Our findings support a key role for butyrate as a dietary metabolite can influence integral transcriptional networks at distinct developmental stages, such as Notch, Wnt and YAP-TAZ, that govern ISC fate within the native stem cell niche [23] and in more mature GLP-1 L-cells to divert to a PYY-biased L-cell identity [43,54].

The FFAR2-selective allosteric modulator, AZ-1729, has been previously characterized as a biased agonist and positive allosteric modulator of FFAR2 activity for only propionate mediated actions [31,67]. Our study also reports the ability of this ligand to act as a PAM for acetate and butyrate at a second messenger level, and that it’s G⍺I signal activity is internalization- independent, suggesting ligand and location bias for AZ-1729. It’s PAM activity enhanced butyrate-induced *Pyy* and *Pax6* expression levels, further supporting a plasma membrane FFAR2/Gαi signal pathway in promoting mature PYY positive L-cells. While employed to date as a pharmacological tool to probe the pleiotropic physiological actions of FFAR2, it also highlights the potential for such ligands to selectively amplify, or reduce, specific SCFA actions, including promoting the abundance of mature PYY L cells from the intestinal stem cell niche or at a later stage of L-cell maturation to modify anorectic gut hormonal content.

Overall, we demonstrate that butyrate shapes distinct transcriptional, endocrine and morphological profiles of enteroendocrine L-cells, and that FFAR2/G⍺i activity is both spatially controlled and has the potential to be selectively amplified via allosteric modulators. Despite the accelerated therapeutic application of GLP-1R agonists, approaches that could harness or modulate endogenous anorectic gut hormone responses in response to diet-induced SCFA production in a selective manner could represent efficacious anti-obesity avenues with improved gastrointestinal function.

## Limitations

While focusing the studies on human L-cell models afforded the availability of FFA2-selelctive antagonists, this pharmacological tool could not be applied in the Hes-1 GFP mouse model as it does not have significant affinity for mouse FFAR2 [68], thus the role of butyrate in regulating NOTCH activity may be via FFAR2 dependent and/or independent mechanisms. In addition, due to lack of sensitive and selective anti-FFAR2 antibodies, a common challenge across the GPCR superfamily, analysis of receptor trafficking required expression of a FLAG-tagged receptor. Thus, while we have demonstrated a role for spatially-regulated FFAR2 signalling from endogenously expressed receptor in NCI-H716 cells, we acknowledge that interpretation of endocytic profiles of human FFAR2 are from the exogenously expressed receptor.

## Author Contributions

Conceptualization, A.H, A.C.H and G.F.; Investigation, A.H.; Resources, C-W.C, A.C.H. and G.F.; Writing – Original Draft, A.H and A.C.H; Writing – Review and Editing, all authors; Visualization, A.H.; Supervision, A.C.H. and G.F.; Funding Acquisition, all authors.

## Acknowledgements

This study was supported by grants from the Biological Sciences Research Council (BBSRC) and the Genesis Research Trust. We thank Professor Kevin Murphy (Imperial College London) for providing NCI-H716 cells for generation of a master cell bank, and Stephen Rothery at the Facility for Imaging of Light Microscopy (FILM), Imperial College London, for technical assistance with super-resolution confocal microscopy. Biorender was used for the creation of the graphical abstract and schematics.

## STAR Methods

### KEY RESOURCES TABLE

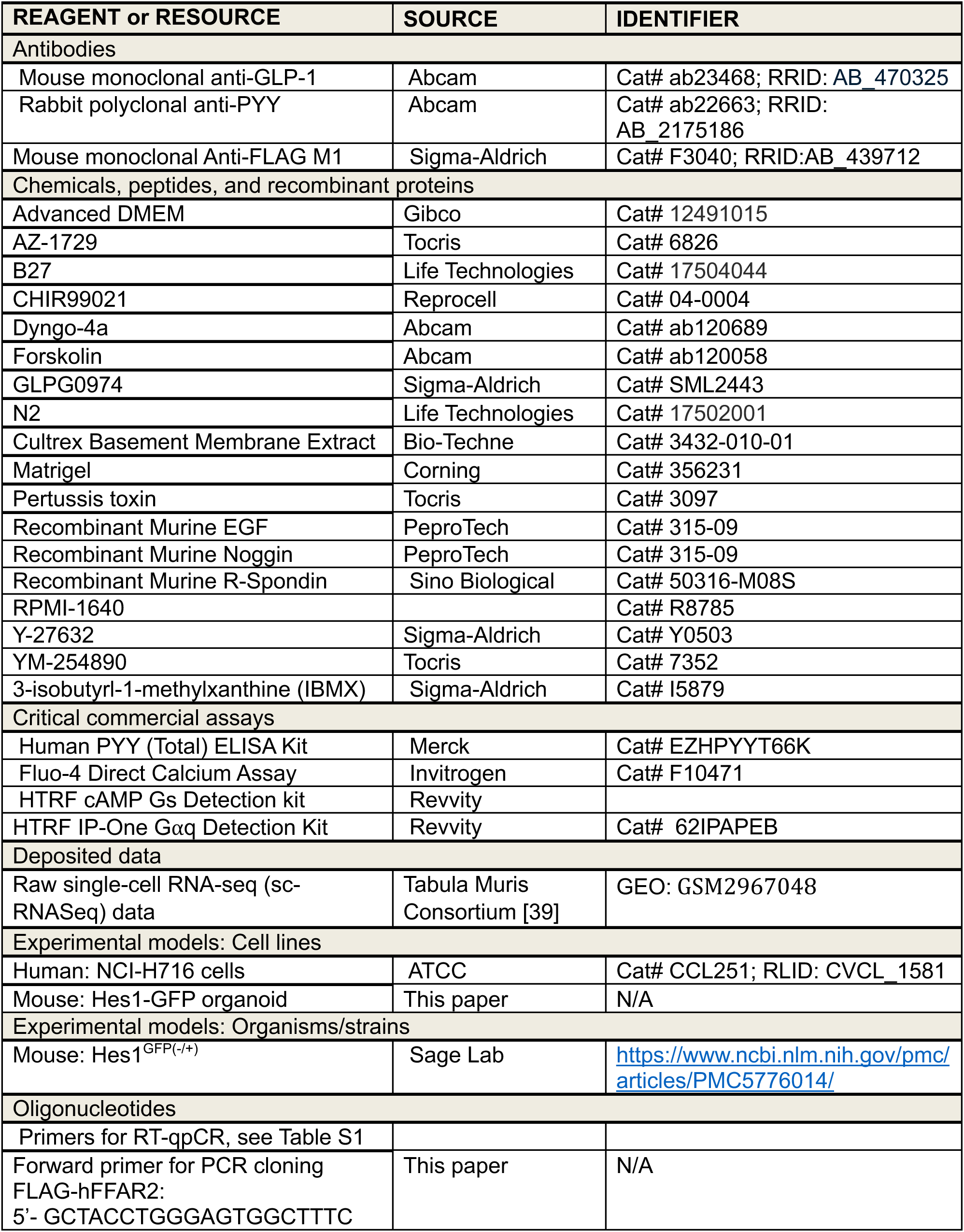
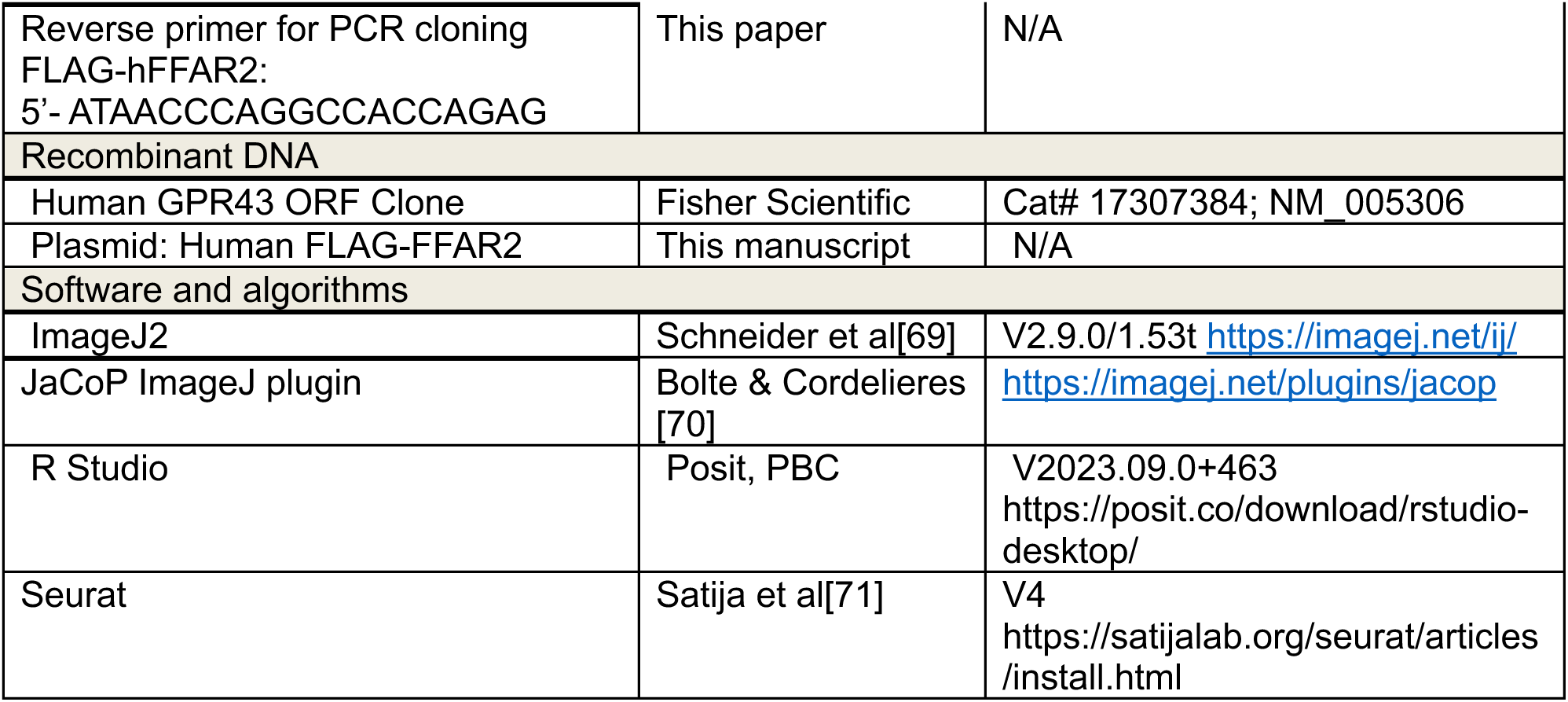

### RESOURCE AVAILABILITY

#### Lead Contact

Further information and requests for resources and reagents should be directed to and will be fulfilled by the lead contact, Aylin Hanyaloglu.

#### Materials Availability

FLAG-hFFAR2 construct is available from the lead contact under a materials transfer agreement with Imperial College London.

### EXPERIMENTAL MODEL AND SUBJECT DETAILS

#### NCI-H716 cell line

NCI-H716 cells (ATCC, CCL-251) were maintained in suspension in RPMI-1640 (Sigma-Aldrich) containing 2g/L D-glucose and 2mM glutamine supplemented with 10% (v/v) fetal bovine serum (Merck) and 100U/ml penicillin/streptomycin (Sigma-Aldrich). in a humidified incubator at 37°C with 5% CO2. Two days before experiments, cells were seeded onto culture plates were pre-coated with Cultrex Basement Membrane Extract (170μl/cm^2^) (R&D Systems) diluted to 0.5mg/ml.

#### Hes1-GFP mice & organoids

Organoids were established from Hes1-GFP reporter mice [72] aged 19-34 weeks. Mice were sacrificed by euthanasia with carbon dioxide asphyxiation, and death was ensured by cervical dislocation. Organoids were maintained in complete crypt culture medium, described in ‘Crypt isolation & culture’, or upto five passages in a humidified incubator with 5% CO_2_ at 37°C, with routine passaging by mechanical dissociation every 7-10 days.

### METHOD DETAILS

#### Crypt isolation & culture

Colons from Hes1-GFP mice were removed, flushed with ice-cold PBS and opened longitudinally [73]. After discarding the distal colon, the proximal colon was cut into 0.5cm fragments and incubated with ice-cold PBS containing 10mM EDTA at 4°C for 45 minutes with rocking. Tissues were cleaned thoroughly with PBS and crypts were mechanically released by vigorous shaking. Isolated crypts were pelleted by centrifugation at 300 x g for 5 min at 4°C. The pellet was resuspended in Matrigel (growth factor reduced; Corning) and crypt media, and then seeded as 25ul domes onto flat bottom 48-well plates (Corning) which were left to solidify for 20 minutes at 37°C. 250μl of crypt media was then added on top of domes and changed every three days. Crypt media consisted of Advanced DMEM (Gibco) supplemented with 1% Glutamax (Gibco) and 1% Penicillin-Streptomycin (Gibco) along with the following components: 40ng/ml Recombinant EGF (Peptrotech), 200ng/ml Recombinant Noggin (Peprotech) and 500ng/ml Recombinant R-spondin (Sino Bioscience), 1x B27 supplement (Life Technologies), 1x N2 supplement (Life Technologies), 3μM CHIR99021 (LC laboratories), and 10μM Y-27632 dihydrochloride monohydrate (10μM, Sigma-Aldrich). Cultures were per maintained in a humidified incubator with 5% CO2 at 37°C.

#### Reagents & antibodies

Antibodies used were mouse anti-GLP-1 (Abcam); rabbit anti-PYY (Abcam) and mouse anti-FLAG M1 (Sigma-Aldrich). Acetate, propionate and butyrate were used as sodium salts (Sigma-Aldrich) and 100mM stocks were made up fresh in PBS. Other agonists used were AZ-1729 (Tocris) at 1µM, and forskolin (Abcam) at 3µM. The inhibitors used were 3-Isobutyl-1- methylxanthine (IBMX) (Sigma-Aldrich) at 0.5mM (5 min pre-treatment), GLPG0974 (Sigma-Aldrich) at 1µM (15 min pre-treatment), Pertussis toxin (PTX) (Tocris) at 500ng/ml (20 hour pre-treatment), YM-254890 (Tocris) at 10µM (15 min pre-treatment) and Dyngo-4a (Abcam) at 50mM (45 min pre-treatment).

#### Plasmid construction & transfection

FLAG-tagged human FFAR2 (FLAG-hFFAR2) was generated by PCR amplification of FFAR2 from a human FFAR2 plasmid (Origene), and ligation at EcoRV and AfeI sites of a FLAG LHR/pcDNA3.1 plasmid via digestion. FLAG-hFFAR2 was transfected into NCI-H716 cells during the seeding process as follows. FLAG-hFFAR2 was incubated with Lipofectamine 2000 (Invitrogen) for complex formation, followed by direct addition of the complex into the cellular suspension. The cells were then seeded in RPMI medium supplemented with 10% fetal bovine serum (FBS) without antibiotics and incubated for 48 hours.

#### RNA isolation & Real-Time Quantitative PCR (RT-qPCR)

For RNA isolation, NCI-H716 cells were seeded in 6 well plates at a density of 1.2 – 1.5 X 10^6^ cells/well, and RNA was isolated using TRIzol reagent (Invitrogen) as per manufacturer’s protocol. Extracted RNA was treated with DNAse I (Invitrogen) to ensure removal of contaminant genomic DNA, and 1µg was reverse transcribed to cDNA with SuperScript IV Reverse Transcriptase (Invitrogen). Target-specific primers were designed manually using National Center for Biotechnology Information Primer-BLAST Tool. Target genes were amplified (*Gcg, Pyy, Ffar2, Ffar3, Atoh1, Ngn3 NeuroD1*, *Pax4, Arx Pax6*), alongside *Rpl12* housekeeping gene with custom primers **(Table S1)**, 2x PowerUp SYBR Master Mix (ThermoFisher) and nuclease-free water in a StepOne Plus Real-Time PCR system (Applied Biosystems). PCR conditions consisted of an initial biphasic incubation at 50°C at 5 min and then 95°C for 5 min, followed by 40 cycles of 15 secs at 95°C and 1 min at 65°C. qPCR Samples were run in at least triplicate. PCR product specificity was confirmed by post-PCR melt-curve analysis which consisted of the following conditions: 95°C for 15 seconds, 60°C for 1 min, and then incremental temperature increases by 0.3°C every 15 seconds upto 95°C. The 2^-ΔΔCt^ method was used calculate relative changes in gene expression [74].

#### Microscopy-based Quantitative Assays

##### Fixed sample preparation

Confocal images of fixed samples were acquired using a Leica Stellaris 8 Inverted confocal microscope (Leica) with the LasX software. NCI-H716 were plated on 13mm coverslips in 24- well plates at a density of 7.5 x 10^4^/well. For receptor trafficking analyses, FLAG-hFFAR2- transfected cells were stained with FLAG M1 antibody for 30 min at 37°C, with indicated ligands added in for the last 20 min of the incubation. If inhibitors were used, these were administered prior to M1 treatment. M1 antibody was stripped off cell membranes by washing cells three times with 0.04M EDTA in PBS. Next, cells were fixed by incubation with 4% paraformaldehyde in PBS for 30 minutes, blocked in 2% FBS for 1hr, and then permeabilised with 0.2% Triton X-100 in PBS-Ca^2+^ for 15 mins. For gut hormone immunofluorescence analysis, cells were incubated with primary antibody(s) for 1hr at this point and then washed. Next, cells were incubated with goat anti-mouse (Invitrogen) or anti-rabbit (Invitrogen) Alexa secondary antibodies for one hour in the dark, washed, and mounted onto glass slides using Fluoromount-G (Invitrogen). To stain cell cytoplasm in morphology workflows, HCS CellMaskTM Deep Red (Invitrogen) was added during the last 30 min of secondary antibody incubation.

##### Hormone granule quantification

For quantification of GLP-1 and PYY, the LIGHTNING detection module on Leica Stellaris 8 was applied, utilising a pinhole size of 0.5 Airy Units (AU) and adaptive deconvolution to maximize image resolution. Background fluorescence was corrected in FIJI (ImageJ) using the ‘Subtract’ function. Individual cells were manually outlined as regions of interest (ROIs). Channel-specific autothresholds were determined and consistently applied across all treatment conditions. Granules were quantified using the ‘Analyse Particles’ tool, with the lower limit for particle diameter set to 120nm. Measurements were extracted per ROI for downstream analysis.

##### Cellular morphology analysis

For measuring morphological parameters in NCI-H716 cells, automatic particle analysis was performed using FIJI (ImageJ) in cells stained with HCS CellMask™ Deep Red. To minimize observer bias, treatment allocation was blinded during the image analysis process. Binary masks were generated from single-channel images of the CellMask™ stain, with manual thresholding parameters established and maintained consistently across all experimental conditions. In more confluent samples, where cells were in close proximity or touching, the ‘watershed’ function was applied to separate adjacent cells. The ‘Analyze Particles’ function was used to identify regions of interest (ROIs) corresponding to individual cells, with a minimum particle area of 20 µm² set to exclude non-cellular background artifacts. The binary mask was then overlaid onto the original CellMask™ channel to verify the accurate identification of "true" cells by the automated analysis. Quantitative measurements, including area, Feret diameter, and circularity, were extracted for each cell.

##### Live intracellular Ca^2+^ accumulation

Intracellular Ca2+ accumulation in live NCI-H7176 cells was captured at 1s intervals with a SP5 inverted confocal microscope (Leica) with LasX acquisition software. NCI-H716 cells were plated on 35mm dishes (Mattek) with 14mm x 1.5mm glass coverslips at a density of 8 x 10^5^ cells/dish. Cells were incubated with Fluo-4 Direct^TM^ calcium indicator (Invitrogen) as per manufacturer’s instructions for 30 minutes in an incubator at 37°C in the dark, and 30 minutes at room temperature in the dark. Fluorescent intensity was quantified in at least 50 cells per sample from using the Fiji Time series analyser plugin, and subsequently averaged across cells in each sample.

##### Live imaging of Hes1-GFP organoid cultures

Hes1-GFP organoids in culture were imaged using a widefield fluorescence microscope (Keyence). Image analysis was carried out on FIJI, whereby outlines of individual organoids, excluding the lumenal area, were manually assigned as unique ROIs. GFP fluorescence was calculated as the average pixel intensity (i.e mean grey value) per ROI corrected to background. Total area was also computed for individual ROIs, with manual thresholding to background to determine % GFP-negative area.

#### Intracellular cAMP and IP1 signalling assays

NCI-H716 cells were plated in 96-well plates at a density of 6 x 10^4^ cells/well and cAMP and IP1 accumulation were measured by homogeneous time-resolved fluorescence (HTRF). To measure intracellular cAMP accumulation, NCI-H716 cells were pre-treated with 0.5mM 3- isobutyl-1- methylxanthine (IBMX) (Sigma) for 5 min prior to any ligand treatments indicated. For IP1 measurements, NCI-H716 cells were treated with indicated ligands in the presence of 50mM LiCl. Cells were lysed in Phospho-total Lysis buffer 4 (Revvity) and centrifuged at 16,000 x g for 15 min at room temperature. Supernatants were analysed using a HTRF cAMP Gs Dynamic Detection Kit (Revvity) or a HTRF IP-One G⍺q Detection Kit (Revvity) as per manufacturer’s instructions, and PHERAstar Plus reader (BMG LABTECH).

#### PYY secretion assay

NCI-H716 cells were grown on 24 well plates at a density of 5 x 10^5^ cells/well and treated with Krebs buffer containing 0.2% fatty-acid free BSA (Sigma-Aldrich) and indicated ligand(s). Supernatants were collected and centrifuged at 800 x g for 10 mins at 4°C, and PYY concentrations were measured by ELISA (Thermo Fisher) as per manufacturer’s instructions. Cell lysates were prepared using RIPA lysis buffer and centrifuged at 16000 x g for 15 min. Lysate protein content was determined by Coomassie Blue (Bradford) assay (Thermo Fisher) and used to normalise PYY secretion values.

#### Single-cell RNA-seq (scRNA-seq) data analysis

Single-cell RNA-seq data from the Tabula Muris Project, generated by FACS-based full-length transcript analysis, were downloaded from the Gene Expression Omnibus (GEO; accession GSM2967048) [39]. Data were processed using the Seurat package in R [71]. Raw gene count matrices were imported and converted into Seurat objects using the CreateSeuratObject function. Cells were retained for downstream analysis if they expressed at least 500 genes and had a minimum of 50,000 unique molecular identifier (UMI) counts, resulting in a dataset comprising 3,950 large intestinal cells and expression profiles for 23,433 genes. Data normalization and scaling were performed using Seurat’s LogNormalize method. Cell-type annotations—epithelial cells, enterocytes, goblet cells, brush cells, and enteroendocrine cells (EECs)—were assigned by integrating cell ontology metadata provided with the Tabula Muris dataset.

### QUANTIFICATION AND STATISTICAL ANALYSIS

All statistical data analyses were performed on GraphPad Prism 9. When comparing two groups, one-tailed unpaired *t* test was used to determine statistical significance. For multiple comparisons, one-way ANOVA was used with Tukey’s post-hoc test when comparing every mean with every other mean, or Dunnett’s post-hoc test when comparing all means to a control mean. When comparing two groups under multiple conditions, two-way ANOVA with Tukey’s post-hoc test was used. When determining significance of treatment group values normalised to control, one-sample t tests were carried out against a bounded control value (1). For all tests, significance was considered at p < 0.05.

## REFERENCES

1. Waddell, I.S., and Orfila, C. (2023). Dietary fiber in the prevention of obesity and obesity-related chronic diseases: From epidemiological evidence to potential molecular mechanisms. Crit Rev Food Sci Nutr 63, 8752–8767. 10.1080/10408398.2022.2061909.

2. Dagbasi, A., Byrne, C., Blunt, D., Serrano-Contreras, J.I., Franco Becker, G., Blanco, J.M., Camuzeaux, S., Chambers, E., Danckert, N., Edwards, C., et al. (2024). Diet shapes the metabolite profile in the intact human ileum, which affects PYY release.

3. Freeland, K.R., Wilson, C., and Wolever, T.M.S. (2010). Adaptation of colonic fermentation and glucagon-like peptide-1 secretion with increased wheat fibre intake for 1 year in hyperinsulinaemic human subjects. British Journal of Nutrition 103, 82–90. 10.1017/S0007114509991462.

4. Bodnaruc, A.M., Prud’Homme, D., Blanchet, R., and Giroux, I. (2016). Nutritional modulation of endogenous glucagon-like peptide-1 secretion: A review. Nutr Metab (Lond) 13. 10.1186/s12986-016-0153-3.

5. Spreckley, E., and Murphy, K.G. (2015). The L-Cell in Nutritional Sensing and the Regulation of Appetite. Front Nutr 2. 10.3389/fnut.2015.00023.

6. Hickey, J.W., Becker, W.R., Nevins, S.A., Horning, A., Perez, A.E., Zhu, C., Zhu, B., Wei, B., Chiu, R., Chen, D.C., et al. (2023). Organization of the human intestine at single-cell resolution. Nature 619, 572–584. 10.1038/s41586-023-05915-x.

7. Le Poul, E., Loison, C., Struyf, S., Springael, J.Y., Lannoy, V., Decobecq, M.E., Brezillon, S., Dupriez, V., Vassart, G., Van Damme, J., et al. (2003). Functional characterization of human receptors for short chain fatty acids and their role in polymorphonuclear cell activation. Journal of Biological Chemistry 278, 25481–25489. 10.1074/jbc.M301403200.

8. Brown, A.J., Goldsworthy, S.M., Barnes, A.A., Eilert, M.M., Tcheang, L., Daniels, D., Muir, A.I., Wigglesworth, M.J., Kinghorn, I., Fraser, N.J., et al. (2003). The orphan G protein-coupled receptors GPR41 and GPR43 are activated by propionate and other short chain carboxylic acids. Journal of Biological Chemistry 278, 11312–11319. 10.1074/jbc.M211609200.

9. Chambers, E.S., Viardot, A., Psichas, A., Morrison, D.J., Murphy, K.G., Zac-Varghese, S.E.K., MacDougall, K., Preston, T., Tedford, C., Finlayson, G.S., et al. (2015). Effects of targeted delivery of propionate to the human colon on appetite regulation, body weight maintenance and adiposity in overweight adults. Gut 64, 1744–1754. 10.1136/gutjnl-2014-307913.

10. Tolhurst, G., Heffron, H., Lam, Y.S., Parker, H.E., Habib, A.M., Diakogiannaki, E., Cameron, J., Grosse, J., Reimann, F., and Gribble, F.M. (2012). Short-chain fatty acids stimulate glucagon-like peptide-1 secretion via the G-protein-coupled receptor FFAR2. Diabetes 61, 364–371. 10.2337/db11-1019.

11. Psichas, A., Sleeth, M.L., Murphy, K.G., Brooks, L., Bewick, G.A., Hanyaloglu, A.C., Ghatei, M.A., Bloom, S.R., and Frost, G. (2015). The short chain fatty acid propionate stimulates GLP- 1 and PYY secretion via free fatty acid receptor 2 in rodents. Int J Obes 39, 424–429. 10.1038/ijo.2014.153.

12. Caengprasath, N., Gonzalez-Abuin, N., Shchepinova, M., Ma, Y., Inoue, A., Tate, E.W., Frost, G., and Hanyaloglu, A.C. (2020). Internalization-Dependent Free Fatty Acid Receptor 2 Signaling Is Essential for Propionate-Induced Anorectic Gut Hormone Release. iScience 23. 10.1016/j.isci.2020.101449.

13. Sealy, L. (1978). The effect of sodium butyrate on histone modification. Cell 14, 115–121. 10.1016/0092-8674(78)90306-9.

14. Candido, E. (1978). Sodium butyrate inhibits histone deacetylation in cultured cells. Cell 14, 105–113. 10.1016/0092-8674(78)90305-7.

15. Roediger, W.E.W. (1982). Utilization of Nutrients by Isolated Epithelial Cells of the Rat Colon. Gastroenterology 83, 424–429. 10.1016/S0016-5085(82)80339-9.

16. Caengprasath, N., and Hanyaloglu, A.C. (2019). Hardwiring wire-less networks: spatially encoded GPCR signaling in endocrine systems. Curr Opin Cell Biol 57, 77–82. 10.1016/j.ceb.2018.12.009.

17. Sanchez, J.G., Enriquez, J.R., and Wells, J.M. (2022). Enteroendocrine cell differentiation and function in the intestine. Curr Opin Endocrinol Diabetes Obes 29, 169–176. 10.1097/MED.0000000000000709.

18. Richards, P., Pais, R., Habib, A.M., Brighton, C.A., Yeo, G.S.H., Reimann, F., and Gribble, F.M. (2016). High fat diet impairs the function of glucagon-like peptide-1 producing L-cells. Peptides (N.Y.) 77, 21–27. 10.1016/j.peptides.2015.06.006.

19. Petersen, N., Reimann, F., Bartfeld, S., Farin, H.F., Ringnalda, F.C., Vries, R.G.J., Van Den Brink, S., Clevers, H., Gribble, F.M., and De Koning, E.J.P. (2014). Generation of l cells in mouse and human small intestine organoids. Diabetes 63, 410–420. 10.2337/db13-0991.

20. Wölnerhanssen, B.K., Moran, A.W., Burdyga, G., Meyer-Gerspach, A.C., Peterli, R., Manz, M., Thumshirn, M., Daly, K., Beglinger, C., and Shirazi-Beechey, S.P. (2017). Deregulation of transcription factors controlling intestinal epithelial cell differentiation; a predisposing factor for reduced enteroendocrine cell number in morbidly obese individuals. Sci Rep 7, 8174. 10.1038/s41598-017-08487-9.

21. Osinski, C., Le Gléau, L., Poitou, C., de Toro-Martin, J., Genser, L., Fradet, M., Soula, H.A., Leturque, A., Blugeon, C., Jourdren, L., et al. (2021). Type 2 diabetes is associated with impaired jejunal enteroendocrine GLP-1 cell lineage in human obesity. Int J Obes 45, 170–183. 10.1038/s41366-020-00694-1.

22. Apelqvist, Å., Li, H., Sommer, L., Beatus, P., Anderson, D.J., Honjo, T., de Angelis, M.H., Lendahl, U., and Edlund, H. (1999). Notch signalling controls pancreatic cell differentiation. Nature 400, 877–881. 10.1038/23716.

23. Ho, J., Puoplo, N., Pokharel, N., Hirdaramani, A., Hanyaloglu, A.C., and Cheng, C.W. (2024). Nutrigenomic underpinnings of intestinal stem cells in inflammatory bowel disease and colorectal cancer development. Front Genet 15. 10.3389/fgene.2024.1349717.

24. Brooks, L., Viardot, A., Tsakmaki, A., Stolarczyk, E., Howard, J.K., Cani, P.D., Everard, A., Sleeth, M.L., Psichas, A., Anastasovskaj, J., et al. (2017). Fermentable carbohydrate stimulates FFAR2-dependent colonic PYY cell expansion to increase satiety. Mol Metab 6, 48–60. 10.1016/j.molmet.2016.10.011.

25. Larraufie, P., Martin-Gallausiaux, C., Lapaque, N., Dore, J., Gribble, F.M., Reimann, F., and Blottiere, H.M. (2018). SCFAs strongly stimulate PYY production in human enteroendocrine cells. Sci Rep 8. 10.1038/s41598-017-18259-0.

26. Park, J.G., Oie, H.K., Sugarbaker, P.H., Henslee, J.G., Chen, T.R., Johnson, B.E., and Gazdar, A. (1987). Characteristics of cell lines established from human colorectal carcinoma. Cancer Res 47, 6710–8.

27. Reimer, R.A., Darimont, C., Gremlich, S., Rie, V., Tral, N.-M., Rüegg, U.T., Rüegg, R., and Macé, K. (2001). A Human Cellular Model for Studying the Regulation of Glucagon-Like Peptide-1 Secretion.

28. Böttcher, G., Sjölund, K., Ekblad, E., Håkanson, R., Schwartz, T.W., and Sundler, F. (1984). Coexistence of peptide YY and glicentin immunoreactivity in endocrine cells of the gut. Regul Pept 8, 261–266. 10.1016/0167-0115(84)90034-X.

29. El-Salhy, M., Grimelius, L., Wilander, E., Ryberg, B., Terenius, L., Lundberg, J.M., and Tatemoto, K. (1983). Immunocytochemical identification of polypeptide YY (PYY) cells in the human gastrointestinal tract. Histochemistry 77, 15–23. 10.1007/BF00496632.

30. Pizzonero, M., Dupont, S., Babel, M., Beaumont, S., Bienvenu, N., Blanqué, R., Cherel, L., Christophe, T., Crescenzi, B., De Lemos, E., et al. (2014). Discovery and optimization of an azetidine chemical series as a free fatty acid receptor 2 (FFA2) antagonist: From hit to clinic. J Med Chem 57, 10044–10057. 10.1021/jm5012885.

31. Bolognini, D., Moss, C.E., Nilsson, K., Petersson, A.U., Donnelly, I., Sergeev, E., König, G.M., Kostenis, E., Kurowska-Stolarska, M., Miller, A., et al. (2016). A novel allosteric activator of free fatty acid 2 receptor displays unique Gi-functional bias. Journal of Biological Chemistry 291, 18915–18931. 10.1074/jbc.M116.736157.

32. Fre, S., Huyghe, M., Mourikis, P., Robine, S., Louvard, D., and Artavanis-Tsakonas, S. (2005). Notch signals control the fate of immature progenitor cells in the intestine. Nature 435, 964– 968. 10.1038/nature03589.

33. Fre, S., Hannezo, E., Sale, S., Huyghe, M., Lafkas, D., Kissel, H., Louvi, A., Greve, J., Louvard, D., and Artavanis-Tsakonas, S. (2011). Notch Lineages and Activity in Intestinal Stem Cells Determined by a New Set of Knock-In Mice. PLoS One 6, e25785. 10.1371/journal.pone.0025785.

34. Ueo, T., Imayoshi, I., Kobayashi, T., Ohtsuka, T., Seno, H., Nakase, H., Chiba, T., and Kageyama, R. (2012). The role of Hes genes in intestinal development, homeostasis and tumor formation. Development 139, 1071–1082. 10.1242/dev.069070.

35. Dou, Y., Pizarro, T., and Zhou, L. (2022). Organoids as a Model System for Studying Notch Signaling in Intestinal Epithelial Homeostasis and Intestinal Cancer. Am J Pathol 192, 1347– 1357. 10.1016/j.ajpath.2022.06.008.

36. Sato, T., Vries, R.G., Snippert, H.J., Van De Wetering, M., Barker, N., Stange, D.E., Van Es, J.H., Abo, A., Kujala, P., Peters, P.J., et al. (2009). Single Lgr5 stem cells build crypt-villus structures in vitro without a mesenchymal niche. Nature 459, 262–265. 10.1038/nature07935.

37. Sato, T., Stange, D.E., Ferrante, M., Vries, R.G.J., van Es, J.H., van den Brink, S., van Houdt, W.J., Pronk, A., van Gorp, J., Siersema, P.D., et al. (2011). Long-term Expansion of Epithelial Organoids From Human Colon, Adenoma, Adenocarcinoma, and Barrett’s Epithelium. Gastroenterology 141, 1762–1772. 10.1053/j.gastro.2011.07.050.

38. Barker, N., van Es, J.H., Kuipers, J., Kujala, P., van den Born, M., Cozijnsen, M., Haegebarth, A., Korving, J., Begthel, H., Peters, P.J., et al. (2007). Identification of stem cells in small intestine and colon by marker gene Lgr5. Nature 449, 1003–7. 10.1038/nature06196.

39. Schaum, N., Karkanias, J., Neff, N.F., May, A.P., Quake, S.R., Wyss-Coray, T., Darmanis, S., Batson, J., Botvinnik, O., Chen, M.B., et al. (2018). Single-cell transcriptomics of 20 mouse organs creates a Tabula Muris. Nature 562, 367–372. 10.1038/s41586-018-0590-4.

40. Yang, Q., Bermingham, N.A., Finegold, M.J., and Zoghbi, H.Y. (2001). Requirement of Math1 for Secretory Cell Lineage Commitment in the Mouse Intestine. Science (1979) 294, 2155– 2158.

41. Jenny, M., Uhl, C., Roche, C., Duluc, I., Guillermin, V., Guillemot, F., Jensen, J., Kedinger, M., and Gradwohl, G. (2002). Neurogenin3 is differentially required for endocrine cell fate specification in the intestinal and gastric epithelium. EMBO J 21, 6338–6347.

42. Mutoh, H., Fung, B.P., Naya, F.J., Tsai, M.-J., Nishitani, J., and Leiter, A.B. (1997). The basic helix–loop–helix transcription factor BETA2/NeuroD is expressed in mammalian enteroendocrine cells and activates secretin gene expression. Proceedings of the National Academy of Sciences 94, 3560–3564. 10.1073/pnas.94.8.3560.

43. Gehart, H., van Es, J.H., Hamer, K., Beumer, J., Kretzschmar, K., Dekkers, J.F., Rios, A., and Clevers, H. (2019). Identification of Enteroendocrine Regulators by Real-Time Single-Cell Differentiation Mapping. Cell 176, 1158–1173.e16. 10.1016/j.cell.2018.12.029.

44. Hill, M.E., Asa, S.L., and Drucker, D.J. (1999). Essential Requirement for Pax6 in Control of Enteroendocrine Proglucagon Gene Transcription.

45. Beucher, A., Gjernes, E., Collin, C., Courtney, M., Meunier, A., Collombat, P., and Gradwohl, G. (2012). The Homeodomain-Containing Transcription Factors Arx and Pax4 Control Enteroendocrine Subtype Specification in Mice. PLoS One 7, e36449. 10.1371/journal.pone.0036449.

46. Marsich, E., Vetere, A., Di Piazza, M., Tell, G., and Paoletti, S. (2003). The PAX6 gene is activated by the basic helix-loop-helix transcription factor NeuroD/BETA2.

47. Larraufie, P., Doré, J., Lapaque, N., and Blottière, H.M. (2017). TLR ligands and butyrate increase *Pyy* expression through two distinct but inter-regulated pathways. Cell Microbiol 19, e12648. 10.1111/cmi.12648.

48. Martin-Gallausiaux, C., Larraufie, P., Jarry, A., Béguet-Crespel, F., Marinelli, L., Ledue, F., Reimann, F., Blottière, H.M., and Lapaque, N. (2018). Butyrate Produced by Commensal Bacteria Down-Regulates Indolamine 2,3-Dioxygenase 1 (IDO-1) Expression via a Dual Mechanism in Human Intestinal Epithelial Cells. Front Immunol 9. 10.3389/fimmu.2018.02838.

49. Ang, Z., Er, J.Z., Tan, N.S., Lu, J., Liou, Y.-C., Grosse, J., and Ding, J.L. (2016). Human and mouse monocytes display distinct signalling and cytokine profiles upon stimulation with FFAR2/FFAR3 short-chain fatty acid receptor agonists. Sci Rep 6, 34145. 10.1038/srep34145.

50. Nilsson, O., Bilchik, A. J., Goldenring, J. R., Ballantyne, G. H., Adrian, T. E., and Modlin, I. M. (1991). Distribution and Immunocytochemical Colocalization of Peptide YY and Enteroglucagon in Endocrine Cells of the Rabbit Colon*. Endocrinology 129, 139–148. 10.1210/endo-129-1-139.

51. Pironi, L., Stanghellini, V., Miglioli, M., Corinaldesi, R., De Giorgio, R., Ruggeri, E., Tosetti, C., Poggioli, G., MorselliLabate, A.M., Monetti, N., et al. (1993). Fat-induced heal brake in humans: A dose-dependent phenomenon correlated to the plasma levels of peptide YY. Gastroenterology 105, 733–739. 10.1016/0016-5085(93)90890-O.

52. Anini, Y., Fu-Cheng, X., Cuber, J.C., Kervran, A., Chariot, J., and Roz□, C. (1999). Comparison of the postprandial release of peptide YY and proglucagon-derived peptides in the rat. Pfl□gers Archiv European Journal of Physiology 438, 299–306. 10.1007/s004240050913.

53. Gerspach, A.C., Steinert, R.E., Schönenberger, L., Graber-Maier, A., and Beglinger, C. (2011). The role of the gut sweet taste receptor in regulating GLP-1, PYY, and CCK release in humans. American Journal of Physiology-Endocrinology and Metabolism 301, E317–E325. 10.1152/ajpendo.00077.2011.

54. Beumer, J., Artegiani, B., Post, Y., Reimann, F., Gribble, F., Nguyen, T.N., Zeng, H., Van den Born, M., Van Es, J.H., and Clevers, H. (2018). Enteroendocrine cells switch hormone expression along the crypt-to-villus BMP signalling gradient. Nat Cell Biol 20, 909–916. 10.1038/s41556-018-0143-y.

55. Karra, E., Chandarana, K., and Batterham, R.L. (2009). The role of peptide YY in appetite regulation and obesity. In Journal of Physiology (Blackwell Publishing Ltd), pp. 19–25. 10.1113/jphysiol.2008.164269.

56. Mannon, P.J., Kanungo, A., Mannon, R.B., and Ludwig, K.A. (1999). Peptide YY/neuropeptide Y Y1 receptor expression in the epithelium and mucosal nerves of the human colon. Regul Pept 83, 11–19. 10.1016/S0167-0115(99)00035-X.

57. Mannon, P. J., and Mele, J. M. (2000). Peptide YY Y1 receptor activates mitogen-activated protein kinase and proliferation in gut epithelial cells via the epidermal growth factor receptor. Biochemical Journal 350, 655–661. 10.1042/bj3500655.

58. Van-Wehle, T., and Vital, M. (2024). Investigating the response of the butyrate production potential to major fibers in dietary intervention studies. NPJ Biofilms Microbiomes 10, 63. 10.1038/s41522-024-00533-5.

59. Baxter, N.T., Schmidt, A.W., Venkataraman, A., Kim, K.S., Waldron, C., and Schmidt, T.M. (2019). Dynamics of Human Gut Microbiota and Short-Chain Fatty Acids in Response to Dietary Interventions with Three Fermentable Fibers. mBio 10. 10.1128/mBio.02566-18.

60. Mayorga-Ramos, A., Barba-Ostria, C., Simancas-Racines, D., and Guamán, L.P. (2022). Protective role of butyrate in obesity and diabetes: New insights. Front Nutr 9. 10.3389/fnut.2022.1067647.

61. Bohórquez, D. V., Chandra, R., Samsa, L.A., Vigna, S.R., and Liddle, R.A. (2011). Characterization of basal pseudopod-like processes in ileal and colonic PYY cells. J Mol Histol 42, 3–13. 10.1007/s10735-010-9302-6.

62. Kaelberer, M.M., Rupprecht, L.E., Liu, W.W., Weng, P., and Bohórquez, D. V. (2020). Neuropod Cells: The Emerging Biology of Gut-Brain Sensory Transduction. Annu Rev Neurosci 43, 337–353. 10.1146/annurev-neuro-091619-022657.

63. Bohórquez, D. V., Samsa, L.A., Roholt, A., Medicetty, S., Chandra, R., and Liddle, R.A. (2014). An Enteroendocrine Cell – Enteric Glia Connection Revealed by 3D Electron Microscopy. PLoS One 9, e89881. 10.1371/journal.pone.0089881.

64. Kaelberer, M.M., Buchanan, K.L., Klein, M.E., Barth, B.B., Montoya, M.M., Shen, X., and Bohórquez, D. V. (2018). A gut-brain neural circuit for nutrient sensory transduction. Science (1979) 361. 10.1126/science.aat5236.

65. Bohórquez, D. V., Shahid, R.A., Erdmann, A., Kreger, A.M., Wang, Y., Calakos, N., Wang, F., and Liddle, R.A. (2015). Neuroepithelial circuit formed by innervation of sensory enteroendocrine cells. Journal of Clinical Investigation 125, 782–786. 10.1172/JCI78361.

66. Thangaraju, M., Cresci, G.A., Liu, K., Ananth, S., Gnanaprakasam, J.P., Browning, D.D., Mellinger, J.D., Smith, S.B., Digby, G.J., Lambert, N.A., et al. (2009). GPR109A Is a G- protein–Coupled Receptor for the Bacterial Fermentation Product Butyrate and Functions as a Tumor Suppressor in Colon. Cancer Res 69, 2826–2832. 10.1158/0008-5472.CAN-08-4466.

67. Lind, S., Holdfeldt, A., Mårtensson, J., Sundqvist, M., Kenakin, T.P., Björkman, L., Forsman, H., and Dahlgren, C. (2020). Interdependent allosteric free fatty acid receptor 2 modulators synergistically induce functional selective activation and desensitization in neutrophils. Biochimica et Biophysica Acta (BBA) - Molecular Cell Research 1867, 118689. 10.1016/j.bbamcr.2020.118689.

68. Sergeev, E., Hansen, A.H., Bolognini, D., Kawakami, K., Kishi, T., Aoki, J., Ulven, T., Inoue, A., Hudson, B.D., and Milligan, G. (2017). A single extracellular amino acid in Free Fatty Acid Receptor 2 defines antagonist species selectivity and G protein selection bias. Sci Rep 7, 13741. 10.1038/s41598-017-14096-3.

69. Schneider, C.A., Rasband, W.S., and Eliceiri, K.W. (2012). NIH Image to ImageJ: 25 years of image analysis. Nat Methods 9, 671–675. 10.1038/nmeth.2089.

70. Bolte, S., and Cordelieres, F.P. (2006). A guided tour into subcellular colocalization analysis in light microscopy. J Microsc 224, 213–232. 10.1111/j.1365-2818.2006.01706.x.

71. Satija, R., Farrell, J.A., Gennert, D., Schier, A.F., and Regev, A. (2015). Spatial reconstruction of single-cell gene expression data. Nat Biotechnol 33, 495–502. 10.1038/nbt.3192.

72. Lim, J.S., Ibaseta, A., Fischer, M.M., Cancilla, B., O’Young, G., Cristea, S., Luca, V.C., Yang, D., Jahchan, N.S., Hamard, C., et al. (2017). Intratumoural heterogeneity generated by Notch signalling promotes small-cell lung cancer. Nature 545, 360–364. 10.1038/nature22323.

73. Cheng, C.W., Biton, M., Haber, A.L., Gunduz, N., Eng, G., Gaynor, L.T., Tripathi, S., Calibasi-Kocal, G., Rickelt, S., Butty, V.L., et al. (2019). Ketone Body Signaling Mediates Intestinal Stem Cell Homeostasis and Adaptation to Diet. Cell 178, 1115–1131.e15. 10.1016/j.cell.2019.07.048.

74. Livak, K.J., and Schmittgen, T.D. (2001). Analysis of Relative Gene Expression Data Using Real-Time Quantitative PCR and the 2−ΔΔCT Method. Methods 25, 402–408. 10.1006/meth.2001.1262.

