## Supplemental Table for "A selective and augmentable FFAR2 signal circuitry programs a butyrate-induced cellular identity of enteroendocrine L-cells"

**Table S1 List of primers used in this study, related to STAR Methods**

**Primers for used for Real-Time Quantitative PCR (RT-qPCR)**

| **Gene** | **Forward** | **Reverse** |
|---|---|---|
| *Ffar2* | 5’- GCTACCTGGGAGTGGCTTTC | 5’- CATAACCCAGGCCACCAGAG |
| *Ffar3* | 5’- CCAATGGGACCTGCTACCTG | 5’- ATCAGCGGGACCACAAAGAG |
| *Gcg* | 5’- TTCAGACCAAAATCACTGAC | 5’- AACATTTCAAACATCCCACG |
| *Pyy* | 5’- TCTTTTCCCATACCGCTGCC | 5’- CTCGTCTGCTTCACAAGCTATC |
| *Atoh1* | 5’- CCTTCCAGCAAACAGGTGAAT | 5’- TTGTTGAACGACGGGATAACAT |
| *Neurog3* | 5’- GCGCCGGTAGAAAGGATGA | 5’- GGTCACTTCGTCTTCCGAGG |
| *Arx* | 5’- TGGAAACAGAGGACCAGCAC | 5’- GTTGGAGTTGGAGCGAGGTT |
| *Pax4* | 5’- ATACCCGGCAGCAGATTGTG | 5’- AAGACACCTGTGCGGTAGTAA |
| *NeuroD1* | 5’- ATGACCAAATCGTACAGCGAG | 5’- GTTCATGGCTTCGAGGTCGT |
| *Pax6* | 5’- CAGACACAGCCCTCACAAACAC | 5’- TGGTGAAGCTGGGCATAGG |
| *Rpl12* | 5’- CATCTCCTTCTCGGCATCA | 5’- AACCCTGTTGTCAATGCCTC |
