## Supplementary Figures for "A selective and augmentable FFAR2 signal circuitry programs a butyrate-induced cellular identity of enteroendocrine L-cells"

**Figure S1. Validation of nutrient-sensing machinery and hormone repertoire characteristic of L-cells in the human colonic NCI-H716 cell line, related to Figure 1**

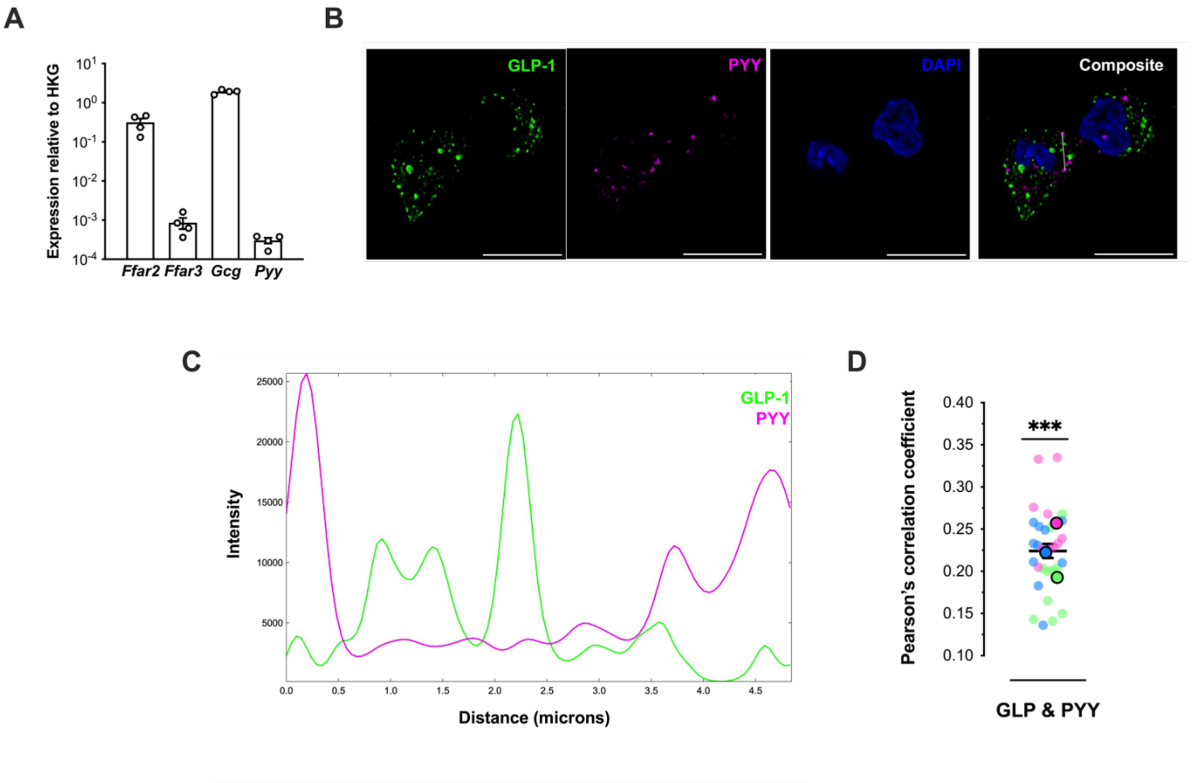


**(A)** *Ffar2, Ffar3, Gcg* and *Pyy* transcript expression in NCI-H716 cells were detected by RT-qPCR. Gene expression relative to ribosomal protein L12 (*Rpl12)* housekeeping gene (HKG) using the $2^{-\Delta Ct}$ method with. Data are presented as the mean ± SEM of data collected across four independent experiments in which conditions were run in at least triplicate. Symbols depict the mean ± SEM of individual biological repeats. **(B-D)** NCI-H716 cells were stained with anti-GLP-1 and anti-PYY antibodies captured by confocal microscopy in super-resolution by an adaptive deconvolution module (LIGHTNING, Leica). **(B)** Representative confocal immunofluorescent images of anti-GLP-1 and anti-PYY staining in NCI-H716 cells Scale bar= 10μM. Images are representative of 40 cells imaged across 6 independent experiments**.** **(C)** Line plot of GLP-1 and PYY intensity profiles across a ROI (indicated by a white line in (A) of 5μM length) **(D)** Pearson's correlation coefficient of GLP-1 and PYY channel overlap in confocal images. Data represented as mean ± SEM of n=30 cells collected across three independent experiments. Data from Independent experiments are colour-coded (pink, blue, green). One sample t-test; Pearsons vs 1 (***p<0.001).

**Figure S2 Butyrate-mediated Gi signalling is inhibited by pertussis toxin and impact of FFAR2 antagonism, G-protein inhibitors, or inhibition of dynamin-mediated internalization on PYY secretion in NCI-H716 cells, related to Figure 3.**

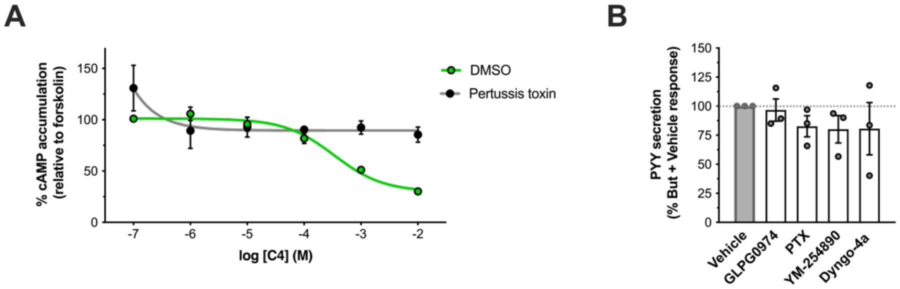


**(A)** Intracellular cAMP accumulation measured in NCI-H716 cells pre-treated with DMSO (vehicle) or Pertussis toxin (500ng/ml, 20 hours) and then stimulated with 3-isobutyl-1- methylxanthine (IBMX) for (500nM,5 min) followed by forskolin (FSK) (3uM) in the presence of increasing doses of But for 5 min. Data are expressed as % response of cAMP accumulation in FSK-treated cells and represent the mean ± SEM of three independent experiments in which conditions were run in triplicates. **(B)** Secretion of PYY from NCI-H716 cells pre-treated with DMSO (Vehicle), GLPG0974 (1uM, 15 min), PTX (500ng/ml, 20 hours), Gaq inhibitor YM-254890 (10nM, 15 min) or dynamin-dependent endocytosis inhibitor Dyngo-4a (50uM, 45 min) followed by incubation with 2mM But for 24h. Cells were incubated with ligand-free secretion buffer for 2hr and supernatants were subsequently assayed for PYY concentration by ELISA. Data was normalised as fold change over untreated cells, and in (D), normalised But with Inhibitor response are shown as % (depicted by dotted line at 100%). Data are presented as the mean ± SEM of data collected across three independent experiments in which conditions were run in at least triplicate. Symbols depict the mean ± SEM of individual biological repeats.

**Figure S3 Positive allosteric modulation of SCFA-mediated FFAR2-G$\alpha$i signalling by AZ-1729 in NCI-H716 cells, related to Figure 4.**

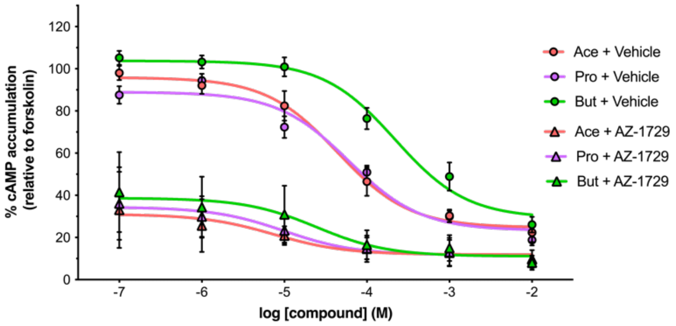


Positive allosteric agonism of SCFA- mediated Gai responses in NCI-H716 cells by AZ-1729. Cells were treated with IBMX (500nM, 5 min), followed by forskolin (FSK) (3μM, 5 min) or combination of FSK (3μM, 5min) and NaCl/SCFA (1mM, 5 min) and DMSO vehicle, or a combination of FSK (3μM, 5min), and NaCl/SCFA (1mM, 5 min), and AZ-1729 (1μM, 5 min). Data represents mean ± SEM (for NaCl) or mean (for SCFAs) of triplicates per independent experiments, n=3 independent experiments.

**Figure S4 Validation of Hes1 expression in the Mouse large intestine and measurements of total Hes1-GFP organoid area in response to butyrate, related to Figure 5**

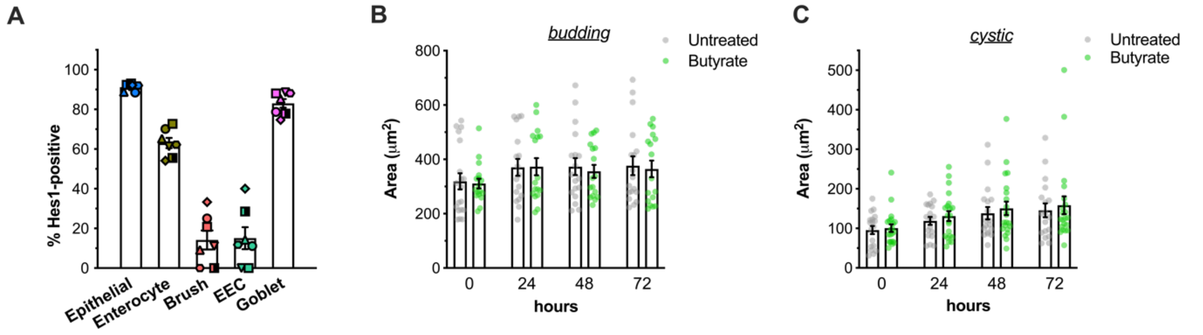


(A) Comparison of Hes1 expression across cellular ontologies in the large intestine, by analysis of the single-cell RNA-seq data from the Tabula Muris project. Data are shown as % of cells per ontology class with log2CPM-normalised Hes1 expression > 0 in Tabula Muris. Distinct symbol shapes represent mean % per cell ontology class in individual mice, n=7 mice**.** Total area of (B) budding and (C) cystic Hes1-GFP organoids monitored across a 72 hour timeframe in untreated culture conditions (Control) or cultures incubated with 2mM of sodium butyrate (But) from 24 hours. Data in are from n=18 'budding' organoids and n=19 'cystic' organoids, and were collected across cultures from three separate Hes1-GFP mice.

**Figure S5 The effect of acetate and propionate on the expression of L-cell transcription factors in NCI-H716 cells, related to Figure 6.**

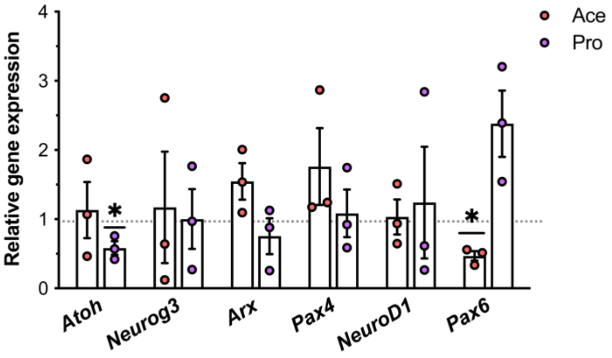


Relative gene expression of *Atoh1, Neurog3, Arx, Pax4, NeuroD1* and *Pax6)* in NCI-H716 cells treated for 24h with 2mM of sodium acetate (Ace) or sodium propionate (Pro). Transcript levels were detected by RT-qPCR and gene expression relative to untreated was determined using the $2^{--\Delta Ct}$ method with ribosomal protein L12 (*Rpl12)* as a housekeeping gene control. Data are presented as the mean ± SEM of data collected across three independent experiments in which conditions were run in at least triplicate. Symbols depict the mean ± SEM of individual biological repeats. (*p<0.05, One sample t-test; Untreated (1) vs SCFA).
